# Determining *Aspergillus fumigatus* transcription factor expression and function during invasion of the mammalian lung

**DOI:** 10.1101/2020.12.23.424128

**Authors:** Hong Liu, Wenjie Xu, Vincent M. Bruno, Quynh T. Phan, Norma V. Solis, Carol A. Woolford, Rachel Ehrlich, Amol C. Shetty, Carie McCraken, Jianfeng Lin, Aaron P. Mitchell, Scott G. Filler

## Abstract

To gain a better understanding of the transcriptional response of *Aspergillus fumigatus* during invasive pulmonary infection, we used a NanoString nCounter to assess the transcript levels of 467 *A. fumigatus* genes during growth in the lungs of immunosuppressed mice. These genes included ones known to respond to diverse environmental conditions and those encoding most transcription factors in the *A. fumigatus* genome. We found that invasive growth *in vivo* induces a unique transcriptional profile as the organism responds to nutrient limitation and attack by host phagocytes. This *in vivo* transcriptional response is largely mimicked by *in vitro* growth in *Aspergillus* minimal medium that is deficient in nitrogen, iron, and/or zinc. From the transcriptional profiling data, we selected 9 transcription factor genes that were either highly expressed or strongly up-regulated during *in vivo* growth. Deletion mutants were constructed for each of these genes and assessed for virulence in mice. Two transcription factor genes were found to be required for maximal virulence. One was *rlmA,* which governs the ability of the organism to proliferate in the lung. The other was *ace1*, which regulates of the expression of multiple secondary metabolite gene clusters and mycotoxin genes independently of *laeA*. Using deletion and overexpression mutants, we determined that the attenuated virulence of the Δ*ace1* mutant is due to decreased expression *aspf1,* which specifies a ribotoxin, but is not mediated by reduced expression of the fumigaclavine gene cluster or the fumagillin-pseruotin supercluster. Thus, *in vivo* transcriptional profiling focused on transcription factors genes provides a facile approach to identifying novel virulence regulators.

**Author summary:** Although *A. fumigatus* causes the majority of cases of invasive aspergillosis, the function of most of the genes in its genome remains unknown. To identify genes encoding transcription factors that may be important for virulence, we used a NanoString nCounter to measure the mRNA levels of *A. fumigatus* transcription factor genes in the lungs of mice with invasive aspergillosis. The transcriptional profiling data indicate that the organism is exposed to nutrient limitation and stress during growth in the lungs, and that it responds by up-regulating genes that encode mycotoxins and secondary metabolites. *In vitro,* this response was most closely mimicked by growth in medium that was deficient in nitrogen, iron and/or zinc. Using the transcriptional profiling data, we identified two transcription factors that govern *A. fumigatus* virulence. These were RlmA, which is governs proliferation in the lung and Ace1, which controls the production of mycotoxins and secondary metabolites.

## Introduction

The fungus *Aspergillus fumigatus* is the major cause of invasive aspergillosis, a progressive pulmonary infection that may disseminate [1–3]. Risk factors for invasive aspergillosis include chemotherapy, corticosteroids, HIV infection, anti-TNF therapy, and solid organ or stem cell transplantation. Because of the growing population of patients at risk of invasive aspergillosis, the annual incidence of this infection has more than tripled since 1990 [2, 4]. Moreover, resistance has emerged to azoles, the front-line therapy for invasive aspergillosis [5, 6]. Therefore, there is an urgent need to understand *A. fumigatus* pathogenicity mechanisms to develop new therapeutic and diagnostic approaches.

Gene expression during infection can provide deep insight into virulence determinants. However, among 10,180 predicted genes in the *A. fumigatus* genome, over 95% are uncharacterized, and fewer than 100 genes have demonstrated roles in virulence. There have been three genome-wide studies of *A. fumigatus* gene expression during *in vivo* infection in the mouse model of pulmonary infection [7–9]. These studies revealed that in early germlings there is up-regulation of respiration, central metabolism, and amino acid biosynthesis genes. At later times, as tissue invasion is initiated, there is up-regulation of cation transport, secondary metabolism, and iron metabolism genes. The authors noted consistent up-regulation of secreted protein genes throughout the infection time-course. These gene expression results are mirrored by functional analysis indicating that defects in iron acquisition, amino acid biosynthesis regulation, and secondary metabolite synthesis all lead to reduced virulence in mouse infection models [1, 10].

These foundational studies investigated gene expression in *A. fumigatus* cells recovered by bronchoalveolar lavage. However, it was not possible to investigate gene expression more than 16 h post-infection because the fungal cells had invaded the lung tissue. To investigate *A. fumigatus* gene expression during tissue invasion, we have taken a different approach, using NanoString technology to assay fungal gene expression in whole lung homogenates. The NanoString nCounter measures RNA levels through probe-based technology and is generally more practical for focused gene set assays than for genome-wide analysis. Here, we used this approach to assay expression of predicted transcription factor genes during invasive growth in the lungs of immunosuppressed mice. Using these data, we selected a panel of transcription factor genes for functional analysis. We found two transcription factor genes, *RlmA* and *Ace1*, with distinct roles in pathogenicity. *RlmA* is required for proliferation in the lung, whereas *Ace1* is required for production of secondary metabolites, especially Asp f1, that mediate pathogenicity. Furthermore, we determined that *in vitro* growth of *A. fumigatus* in *Aspergillus* minimal medium with low zinc and low nitrogen induced a transcriptional response that was similar to response induced during invasive growth in the lung of immunosuppressed mice.

## Results and Discussion

### Invasive infection induces expression of genes involved in nutrient acquisition, stress response, and secondary metabolite biosynthesis

To determine the transcriptional response of *A. fumigatus* during invasive infection, we developed two NanoString probesets. The first was a pilot set that contained probes for 97 genes with functional annotations that included metabolism, iron acquisition, hypoxia, cell wall, and stress response, as described previously [11, 12]. The second contained probes for 400 genes that specify virtually all predicted transcription factors in the *A. fumigatus* genome. We focused on transcription factor genes (TF genes) for three reasons. First, a single transcription factor often controls many functionally related genes, so transcription factor mutants frequently have more prominent phenotypes than mutations in individual target genes [13–17]. Thus, our dataset might be useful for selection of TF genes for functional analysis. Second, there are many methods to identify the indirect or direct targets of a transcription factor, including expression profiling and chromatin immunoprecipitation. Thus, a transcription factor defect can be linked to its target genes to provide physiological and mechanistic insight. Third, transcription factors are prospective drug targets [18], so their analysis can lead to therapeutic benefit.

The transcriptional profiling was performed on RNA isolated from the lungs of mice with invasive aspergillosis, a model that mimics many aspects of human disease [19, 20]. In this model, the mice were immunosuppressed with cortisone acetate and then infected with *A. fumigatus* Af293 via an aerosol chamber, which delivered approximately 5 ×10^3^ conidia to the lungs of each mouse. RNA was prepared from whole lung samples at 2, 4 and 5 days post-infection. In this infection model, mortality begins by day 6 post-infection. There was some overlap between the two sets of probes so the final dataset contained expression data for 467 genes. Data quality was excellent at days 4 and 5 post-infection (S1 Fig); the majority of probes gave detectable signals, and R^2^ values for independent determinations were all >0.95. Data quality was weaker at day 2 post-infection, a reflection of the low *A. fumigatus* titer at that time.

We expected that once *A. fumigatus* achieved steady-state invasive growth, its gene expression profile would be similar in successive time points. Our data support this hypothesis, indicating that there are similar gene expression states at days 4 and 5 post-infection (S1 Table). The mean normalized probe counts for all genes assayed at days 4 and 5 were very similar, with an R^2^ value of 0.96, and no statistical support for differential expression of 88% of genes. The genes that showed most extreme variation between samples were close to the lower detection limits on both days 4 and 5, and thus small changes in the numbers of probe counts lead to large differences in calculated expression ratios. Therefore, our data indicate that gene expression states are very similar on days 4 and 5, shortly before mortality begins.

Growth in the mouse lung resulted in an extensive change in the *A. fumigatus* transcriptional profile compared to growth in *Aspergillus* minimal medium (AMM). Of the 467 genes analyzed, 125 (27%) were up-regulated by at least 2-fold and 85 (18%) were down-regulated by at least 2-fold at 5 days post-infection (S1 Table). There was significant up-regulation of genes involved in iron acquisition, including *fre2*, *hapX*, *sidA*, *sidD, mirB*, and *sit1* (Table 1). Concomitantly, there was down-regulation of *sreA*, whose product represses iron uptake and siderophore synthesis [21]. Other up-regulated genes include *zrfC*, *zrfA*, *aspf2*, and *zafA*, which govern zinc uptake, *nrtB* and *areA*, which control nitrogen uptake, and Afu5g00710, which specifies a GABA permease (Table 1). We infer that growth in the lung imposes limitation for key nutrients, including iron, zinc, and nitrogen, on *A. fumigatus*. Our results are consistent with prior studies of *A. fumigatus* mutants with defects in iron acquisition (Δ*hapX*, Δ*sidA*), zinc homeostasis (Δ*zafA*), and nitrogen uptake (Δ*areA*), all of which have attenuated virulence in mice [22–25].

**Table 1.**
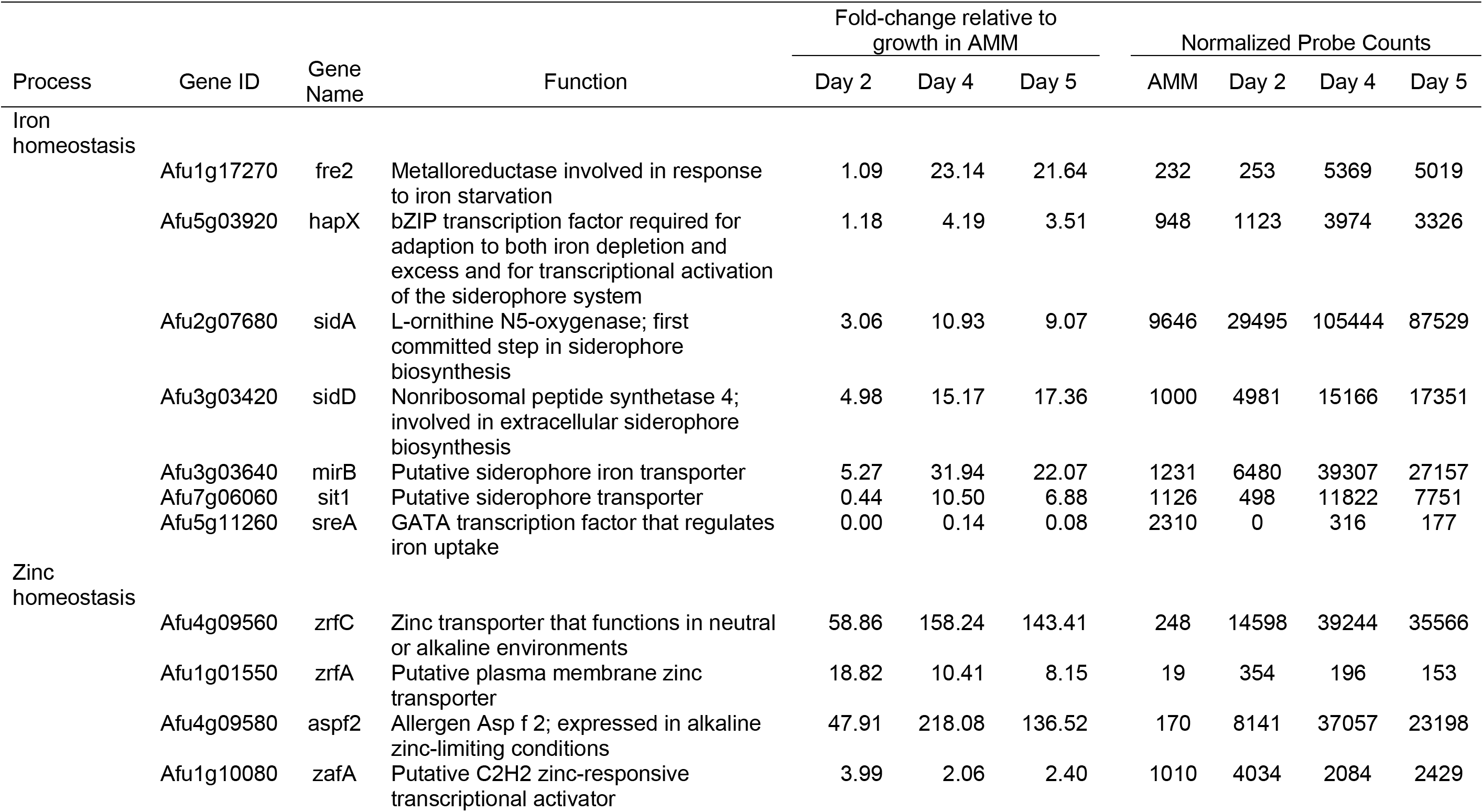
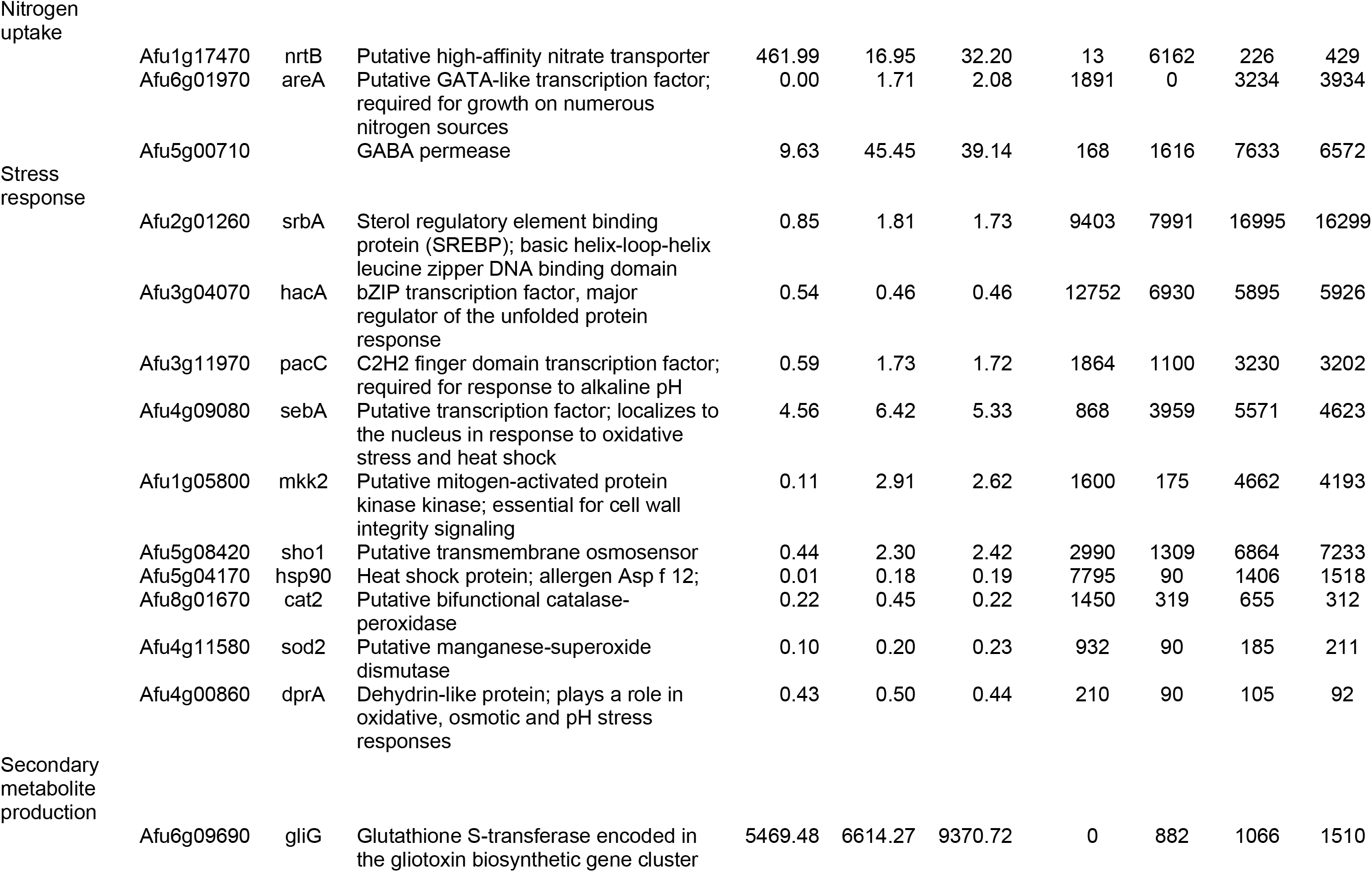
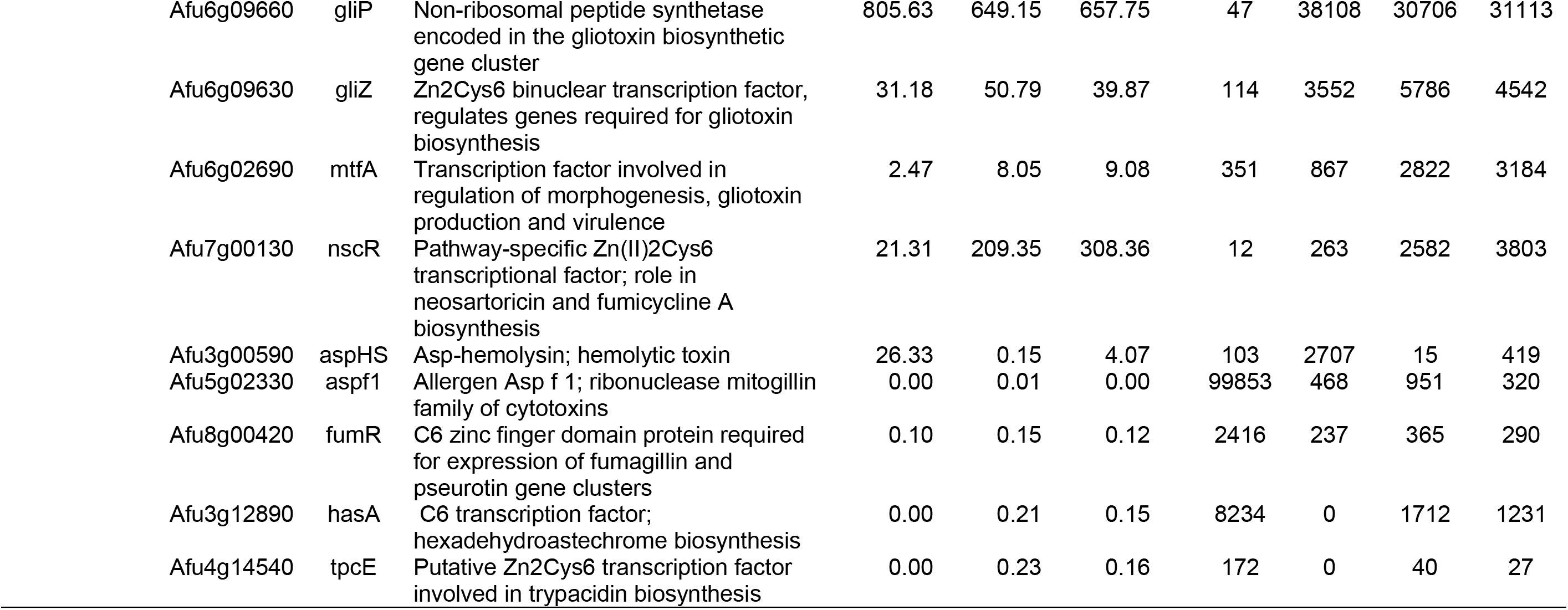
Fold-change and probe counts for selected *A. fumigatus* genes during growth in *Aspergillus* minimal medium (AMM) and during invasive growth in the mouse lung on days 2, 4, and 5.

Some TF genes that are known to govern virulence were not strongly up-regulated by invasive growth in the lung. These include *srbA*, which is required for growth under hypoxia [26], *hacA* which governs the unfolded protein response [27], and *pacC*, which is required for growth under alkaline pH [7]. The expression of these TF genes was increased by less than 2-fold *in vivo* (*srbA* and *pacC*) or even reduced (*hacA*) (Table 1). However, the NanoString counts indicated that all three genes were among the 30 most highly expressed genes *in vivo*. This result suggests that in *A. fumigatus,* TF genes that are highly expressed *in vivo* are likely to govern virulence, even if their expression *in vivo* does not increase relative to growth *in vitro.* We have observed a similar relationship between *in vivo* gene expression levels and virulence regulation in *C. albicans* [13].

In corticosteroid-treated mice with invasive aspergillosis, the invading hyphae stimulate an influx of neutrophils into the lung that accumulate around the organisms [19, 20]. These neutrophils are almost certainly responsible for the up-regulation of genes that are required for normal stress response and virulence in *A. fumigatus*, such as *sebA, mkk2,* and *sho1* [28–30] (Table 1). Surprisingly, some genes involved in stress response were actually down-regulated during in vivo growth. These genes included *hsp90*, *cat2*, *sod2*, and *dprA* (Table 1). Collectively, these results indicate that *in vivo* growth induces expression of only a subset of stress response genes.

Growth in the lung also altered the expression of genes involved in the production of specific secondary metabolites. As compared to organisms grown in AMM, organisms in the mouse lung had a 700-10,000-fold increase in expression of *gliG* and *gliP*, which specify enzymes in the gliotoxin biosynthesis pathway [31–33] (Table 1). Furthermore, there was 40-fold up-regulation of *gliZ*, which encodes the transcription factor that governs gliotoxin synthesis [34], and 9-fold up-regulation of *mtfA*, which induces synthesis of both gliotoxin and extracellular proteases [35] (Table 1). *In vivo* growth also up-regulated expression of *nscR* whose product governs synthesis of neosartoricin [36] and of *aspHS*, which specifies a hemolysin (Table 1). However, *in vivo* growth repressed the expression of *aspf1* which encodes a ribotoxin [12, 37], *fumR*, which governs production of fumagillin and pseurotin [38, 39], *hasA*, which controls production of hexadehydroastechrome [40], and *tpcE*, which regulates production of trypacidin and questin [41] (Table 1). Thus, growth in the mouse lung induces production of a distinct subset of mycotoxins. We speculate that the mycotoxin genes whose expression was down-regulated in the lung must be expressed in other environmental niches, possibly including other anatomic sites within the host.

### Comparisons among gene expression profiles to identify *in vitro* conditions that mimic *in vivo* growth

Similarities among gene expression profiles can reveal parallels among diverse genetic or environmental regulatory inputs. To assess similarity, we adapted our Fisher’s Exact Test (FET) comparison tool for use with *A. fumigatus* gene expression data. We created a database from 129 published microarray or RNA-seq datasets and generated NanoString profiles of *A. fumigatus* during growth in standard AMM and in AMM modified to mimic aspects of *in vivo* growth: limitation for nitrogen, iron, zinc, or oxygen; presence of serum; limitation for combinations of nitrogen, iron, and zinc. Datasets with significant similarity among genes up-regulated on day 5 postinfection (compared to growth in AMM) are listed in Table 2. A heat map clustering of some of these datasets is shown in Figure 1.

**Table 2.** Comparisons among datasets of differentially expressed genes

|  |  |
| --- | --- |
| <u>Dataset</u> | Similarity to lung day 5 vs AMM (P-value) |
| lung day 2 vs AMM | <1.00E-10 |
| lung day 4 vs AMM | <1.00E-10 |
| AMM low N Fe vs AMM | <1.00E-10 |
| AMM low N Fe Zn vs AMM | <1.00E-10 |
| AMM low Zn vs AMM | <1.00E-10 |
| AMM low Fe Zn vs AMM | <1.00E-10 |
| AMM low N Zn vs AMM | <1.00E-10 |
| AMM low Fe vs AMM | 2.38E-10 |
| AMM low N vs AMM | 6.20E-08 |
| lung germling 12-14 hr vs YPD 37° [8] | 4.57E-07 |
| lung germling 12 hr vs spores [7] | 7.30E-07 |
| lung germling 16 hr vs spores [7] | 9.00E-07 |
| AMM+serum vs AMM | 1.35E-06 |
| Description: Genes up-regulated $\geq 2$ fold in our lung day 5 vs AMM dataset were compared to genes up-regulated $\geq 2$ fold in each comparison dataset listed above. The probability that a correlation was due to chance alone was calculated using Fisher's Exact Test. The calculated threshold of significance is a P-value of 0.00017. | |

**Fig 1.**
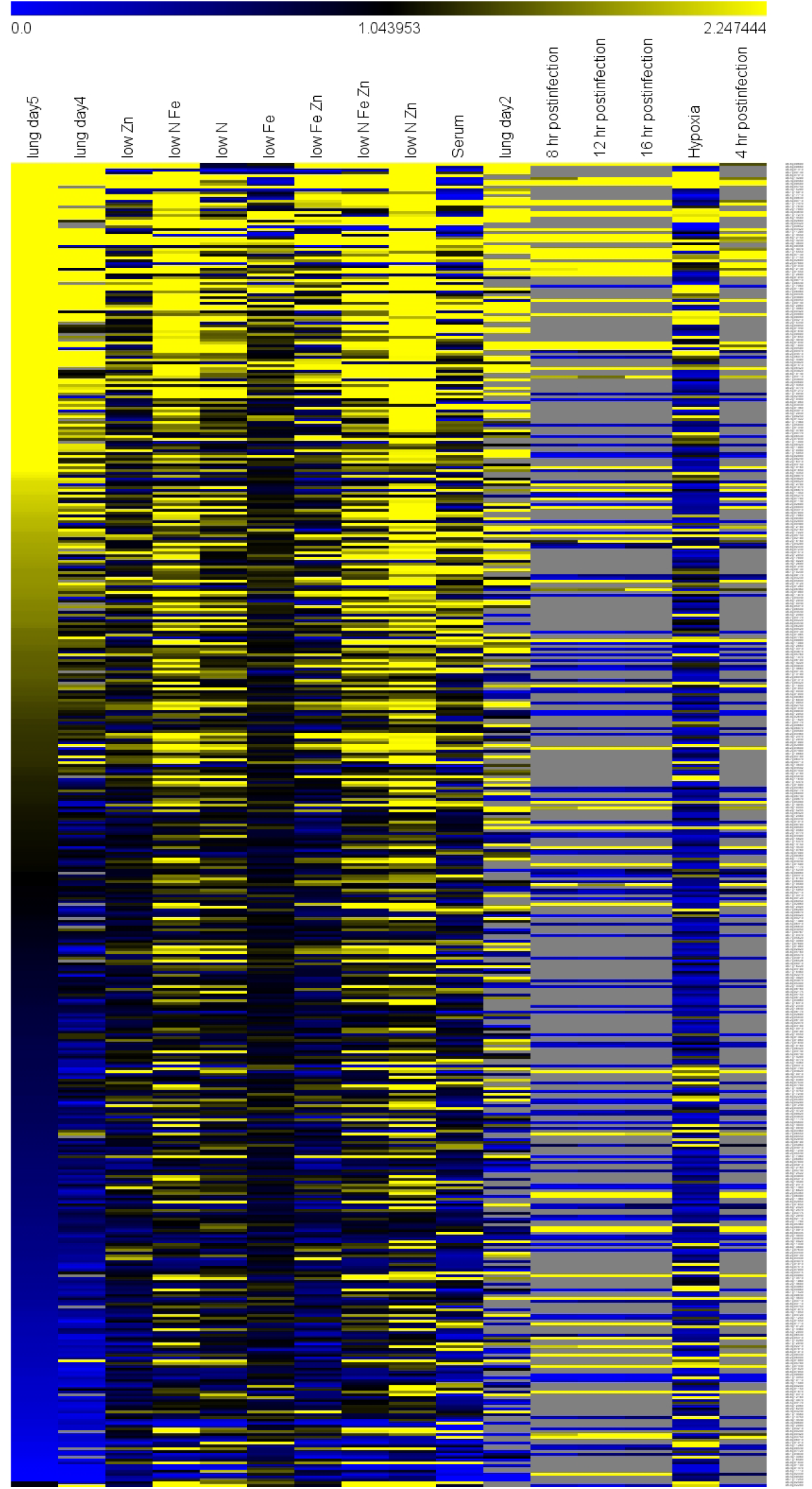
Hierarchical clustering of gene expression datasets. The Nanostring datasets and published datasets were compared by hierarchical clustering based on the 467 genes in the Nanostring datasets. Select dataset are indicated, including lung germlings [7, 8], invasive infection (current data), growth in low zinc or low nitrogen (current *in vitro* data for *Aspergillus* minimal medium (AMM) lacking zinc or nitrate alone, or in combination with each other and with limiting iron), and low iron alone [21]. Grey areas indicated genes with undetectable expression.

These comparisons indicated that our transcriptional profiles of invasive *A. fumigatus* were significantly similar to the published profiles of germlings isolated from the lungs by lavage [7, 8] (Table 2). These correlations make sense, because we would expect common regulatory pathways that govern growth in the lung environment. It is probable that the incomplete overlap between these profiles is due to changes in the microenvironment that the organism experiences as it invades the lung parenchyma and is attacked by phagocytes.

The FET comparison also identified some *in vitro* growth conditions that induced transcriptional profiles that were similar to what was induced by invasive growth in the lungs. While *in vitro* growth in the presence of serum correlated with invasive growth *in vivo,* the most significant correlations were with gene expression in organisms grown in AMM that was limited in nitrogen, iron, zinc, or combinations of the three (Table 2). These correlations also make sense, because all invasive pathogens must combat nutritional immunity imposed by the host. Our data thus fit well with current understanding of *A. fumigatus* infection biology. In addition, these data indicate growing *A. fumigatus* in AMM that is deficient in nitrogen, iron and/or zinc induces a transcriptional response that is quite similar to that induced by invasive growth in the lungs.

### Analysis of deletion mutants to assess transcription factor function in invasive aspergillosis

We sought to use our expression profiling data to prioritize TF genes for functional analysis during invasive aspergillosis. We chose a set of 9 *A. fumigatus* genes that were either highly expressed or highly upregulated during invasive growth in the lung, and whose functions had not been reported previously (Table 3). We created a deletion mutant for each gene and tested the mutants for proliferation in the mouse model of invasive aspergillosis. In this screen, proliferation was assayed by NanoString measurement of *A. fumigatus* rRNA levels relative to mouse housekeeping gene RNA (*ACTB, GAPDH,* and *PPIA*) levels in whole lung homogenates. The Δ*rlmA* mutant displayed 10-fold lower levels of *A. fumigatus* rRNA than the wild-type strain or any other mutant (Fig 2). The Δ*ace1* mutant displayed rRNA levels comparable to those of the wild-type strain, but the lung tissue of the mice infected with the Δ*ace1* mutant appeared to be notably healthier compared to that of mice infected with the wild-type strain. Therefore, we chose to pursue analysis of *rlmA* and *ace1* during infection.

**Table 3.**
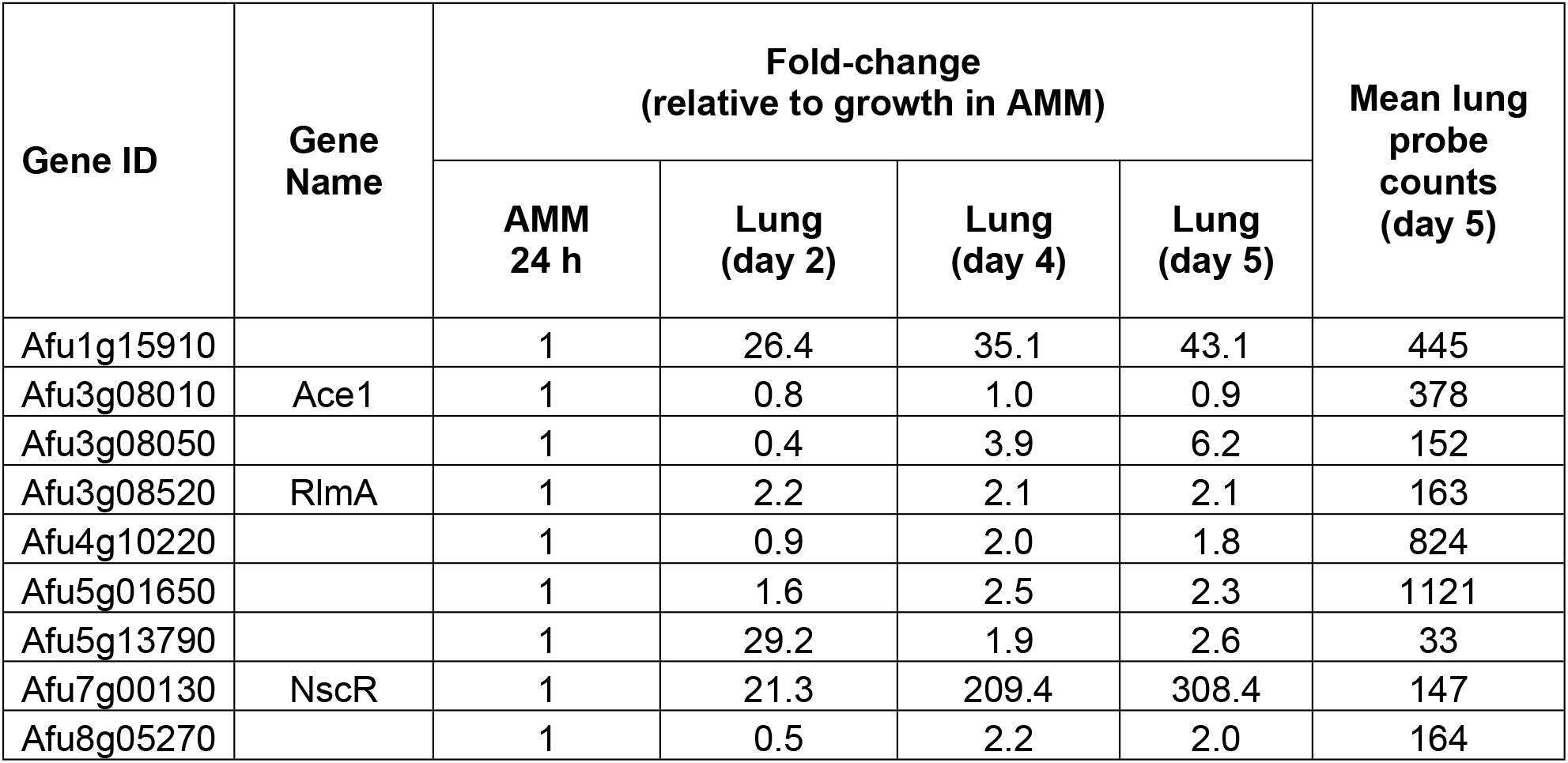
Genes encoding potential transcription factors that were selected for construction of deletion mutants.

**Fig 2.**
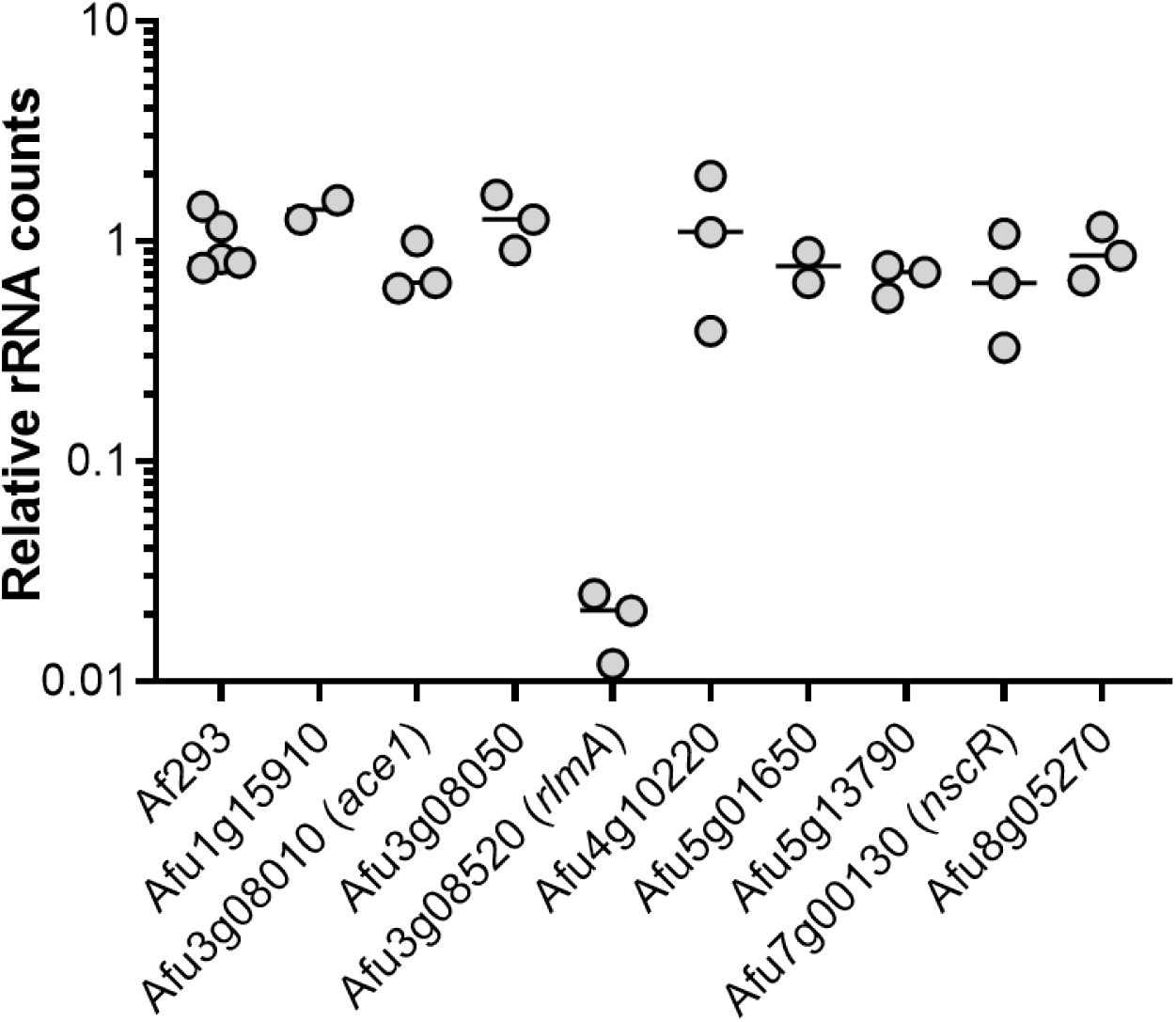
RlmA is required for growth in the lungs during invasive aspergillosis. Pulmonary fungal burden of mice after 5 days of infection with either *A. fumigatus* strain Af293 or mutants deleted for the indicated genes. Lung fungal burden was determined by Nanostring measurement *A. fumigatus* rRNA levels relative to mouse *ACTB, GAPDH,* and *PPIA* levels. Results are from 2-3 mice per strain and are normalized data from mice infected with strain Af293.

### RlmA is required for proliferation in the lung during invasive aspergillosis

RlmA is a putative MADS-box transcription factor whose orthologs in many ascomycetes function in cell wall integrity. *rlmA* RNA levels were down-regulated in the lung germling datasets at 4, 8, and 12 hours post-infection, then began to increase slightly at 16 hours [7]. We found that *rlmA* was up-regulated in our invasive infection datasets at 2, 4, and 5 days post-infection (Table 3). These results led us to the simple hypothesis that RlmA may be required specifically for invasive infection. It is neither up- nor down-regulated 2-fold in any of the *in vitro* published datasets we collected. This observation suggested to us that *rlmA* may be regulated by a signal that is distinctive of the invasive infection environment.

To test RlmA function during invasive aspergillosis, we characterized a Δ*rlmA* deletion mutant in the Af293 background. Mice infected with the Δ*rlmA* mutant survived significantly longer than those infected with the wild-type or Δ*rlmA*+*rlmA* complemented strains (Fig 3A). This result indicates that RlmA is required for proliferation in the lung and pathogenicity in mice immunosuppressed with corticosteroids. Recently, another group reported RlmA is a member of the cell wall integrity pathway in *A. fumigatus* and required for virulence in neutropenic mice [42].

**Fig 3.**
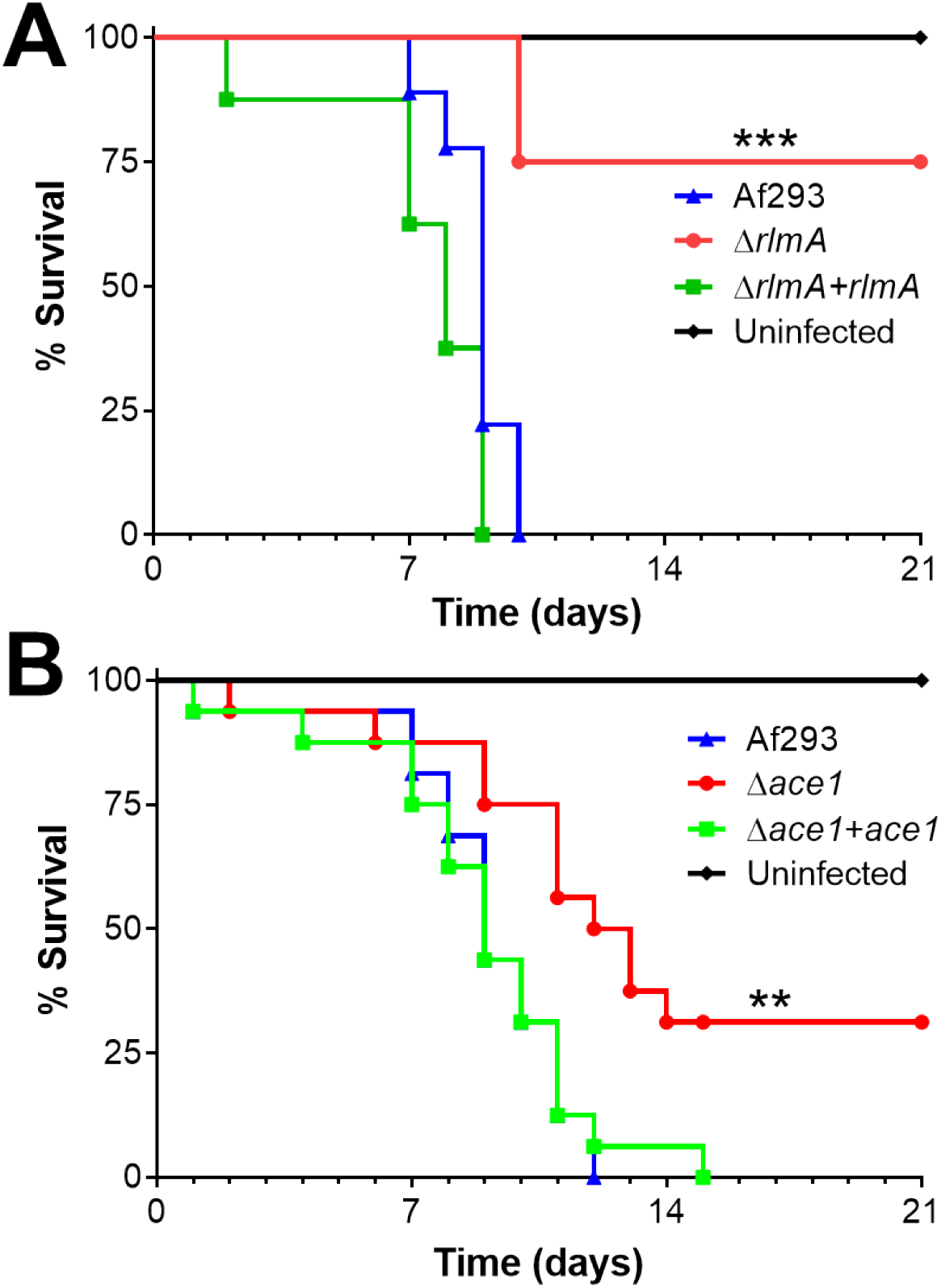
Δ*rlmA* and Δ*ace1* mutants have attenuated virulence. Survival of corticosteroid-immunosuppressed mice infected with the indicated strains. Results are the combined data from two independent experiments, each using 8 mice per strain. **, *p* < 0.01; ***, *p* < 0.001.

### Ace1 governs production of secondary metabolites and toxins during invasive aspergillosis

Ace1 is a C_2_H_2_ zinc finger protein whose *A. nidulans* ortholog, *sltA,* governs ion homeostasis, growth under alkaline pH, and sporulation [43, 44]. We constructed an *A. fumigatus* Δ*ace1* mutant, and found that it grew comparably to the wild-type strain in the presence of high cations and at ph 8 (Supplemental Figure 2). The Δ*ace1* mutant also sporulated similarly to the wild-type strain. Although the Δ*ace1* mutant had increased susceptibility to cell membrane stress caused by protamine and SDS, it had wild-type susceptibility to Congo red, caspofungin, calcofluor white, and hydrogen peroxide protamine (Fig 4). The Δ*ace1* mutant had reduced capacity to adhere to, invade, and damage the A549 pulmonary epithelial cell line, but had wild-type susceptibility to macrophage killing (Fig 5). These results suggest that Ace1 governs the response to cell membrane stress and the capacity of *A. fumigatus* to damage host cells.

**Fig 4.**
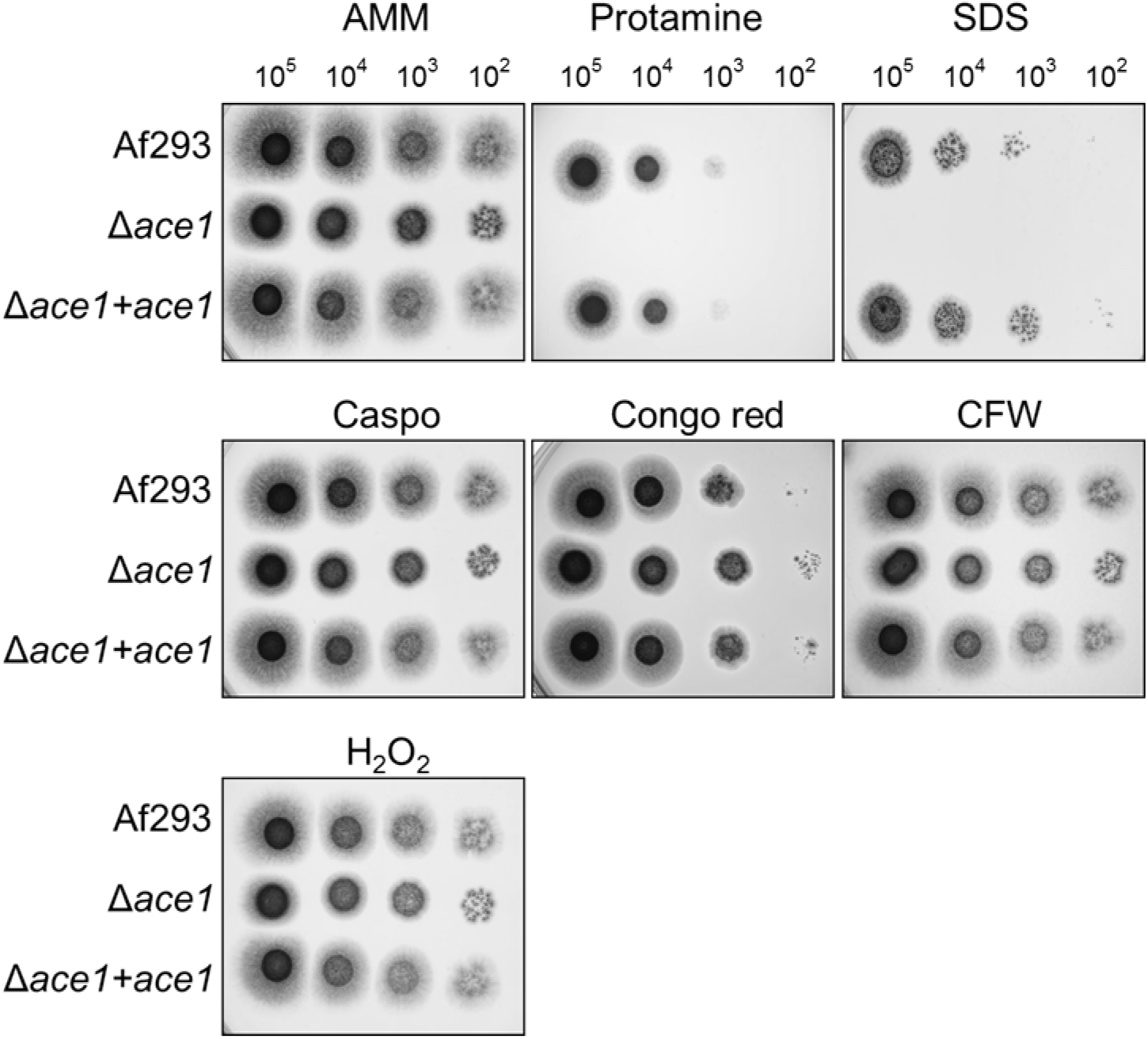
Increased susceptibility of the Δ*ace1* mutant to protamine and SDS. Serial 10-fold dilutions of the indicated strains of *A. fumigatus* were spotted onto *Aspergillus* minimal medium (AMM) containing the indicated stressors. The plates were imaged after incubation at 37°C for 2 d. Caspo, caspofungin; CFW, calcofluor white.

**Fig 5.**
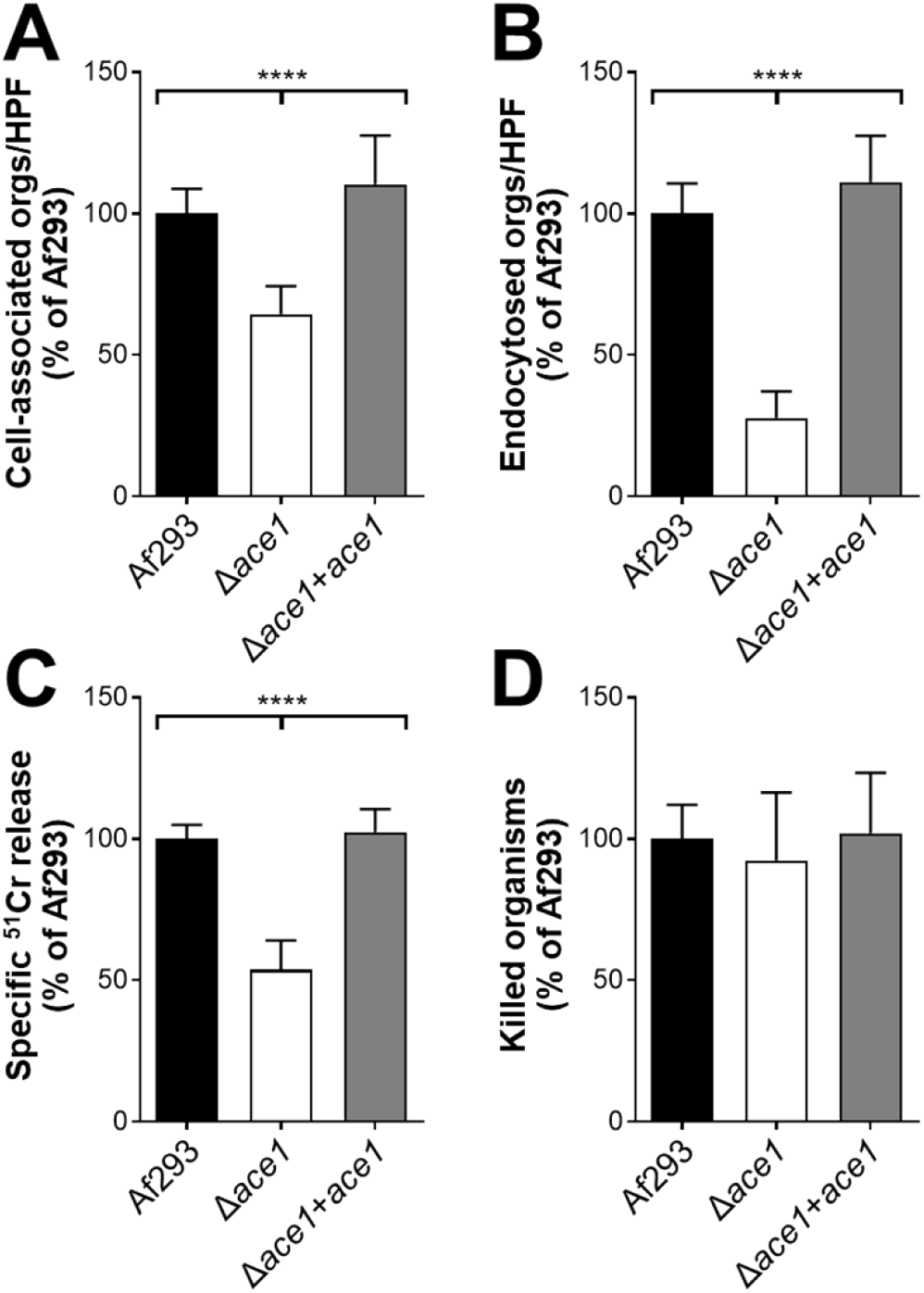
The Δ*ace1* mutant is defective in pulmonary epithelial cell adherence, invasion, and damage. (A-B) The indicated strains of *A. fumigatus* were incubated with the A549 pulmonary epithelial cell line for 2.5 h, after which the number of cell-associated (A; a measure of adherence) and endocytosed (B) organisms was determined by a differential fluorescence assay. (C) The extent of epithelial cell damage induced by the indicated strains after 16 h of infection. (D) The percentage of cells of the indicated *A. fumigatus* strains that were killed by mouse bone marrow-derived macrophages after 8 h of infection. Results are mean ± SD of 3 experiments, each performed in triplicate. Orgs/HPF, organisms per high-powered field; ****, *p* < 0.0001.

In the mouse model of invasive aspergillosis, our rRNA-based titer measurement described above indicated that the Δ*ace1* and wild-type strains proliferated to similar levels in the lung at day 5 post-infection (Fig 2). However, gross inspection of the Δ*ace1* infected lungs suggested that there was less fungal-induced damage. This observation led to the hypothesis that Ace1 may be required for specific pathogenicity functions during invasive aspergillosis rather than for proliferation.

We tested that hypothesis by monitoring mouse survival post-infection in our non-neutropenic invasive aspergillosis model (Figure 3B). We observed that Δ*ace1*-infected mice survived significantly longer than mice infected with the wild-type strain or the Δ*ace1*+*ace1* complemented strain. The finding that mice infected with the Δ*ace1* mutant maintained a high pulmonary fungal burden yet had reduced mortality is similar to what has been found with *A. fumigatus* mutants with defects in secondary metabolite production [12, 31], suggesting that Ace1 may govern the expression of secondary metabolite genes.

To identify Ace1 target genes that might be responsible for the virulence, we performed RNA-seq analysis of the wild-type and Δ*ace1* mutant strains grown in AMM with low nitrogen and low zinc to mimic the conditions during invasive infection in the lung. In *Aspergillus* spp., genes encoding proteins involved in the biosynthesis of secondary metabolites are frequently located in contiguous clusters in the genome. A total of 33 non-overlapping secondary metabolite gene clusters have been identified in *A. fumigatus* Af293 [45]. We found that Ace1 governs the expression of at least 50% of the genes in 16 of these biosynthetic gene clusters (Fig 6, S2 Table). Of these gene clusters, 10 had reduced mRNA expression in the Δ*ace1* mutant, 3 had increased mRNA expression, and 3 had both increased and decreased mRNA expression. In the Δ*ace1* mutant, there was also reduced expression of *aspf1* (Afu5g02330), which encodes a ribotoxin that enhances *A. fumigatus* virulence [12, 37] (S2 Table). We verified the low levels of *aspf1* mRNA in the Δ*ace1* mutant by qPCR (Fig 7A). These results indicate that a principal function of Ace1 is the regulation of production of secondary metabolites and mycotoxins.

**Fig 6.**
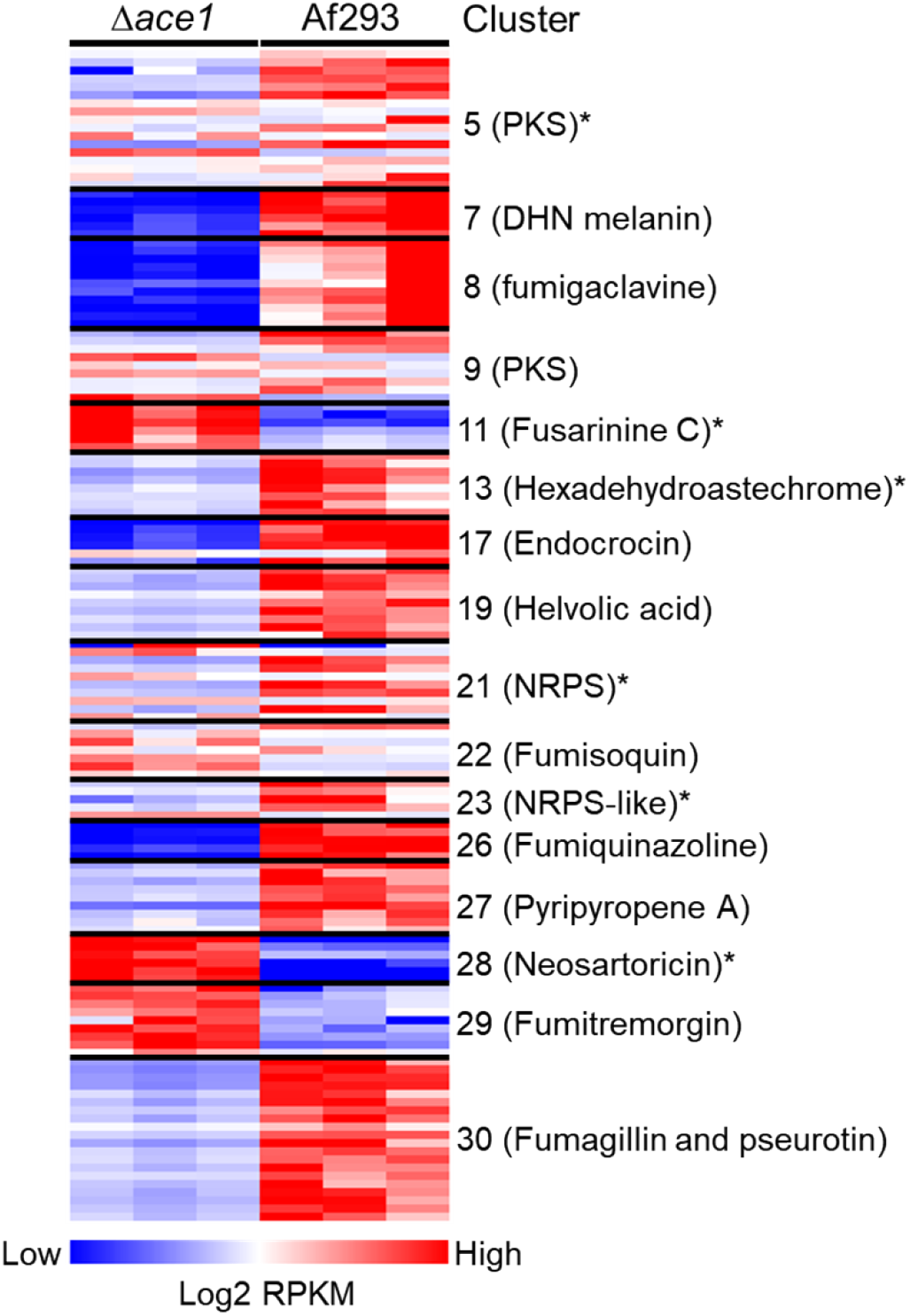
Ace1 governs the expression of secondary metabolite gene clusters. Heat map showing secondary metabolite gene clusters in which the expression of at least 50% of genes were altered in the Δ*ace1* mutant relative to strain Af293. The transcript levels were assessed by RNA-seq analysis of organisms that were grown in liquid AMM with low nitrogen and low zinc in biological triplicate. Secondary metabolite cluster numbers are from [45]. *, gene clusters that were not found to be regulated by LaeA by microarray analysis [46]; NRPS, non-ribosomal peptide synthase; PKS, polyketide synthase.

**Fig 7.**
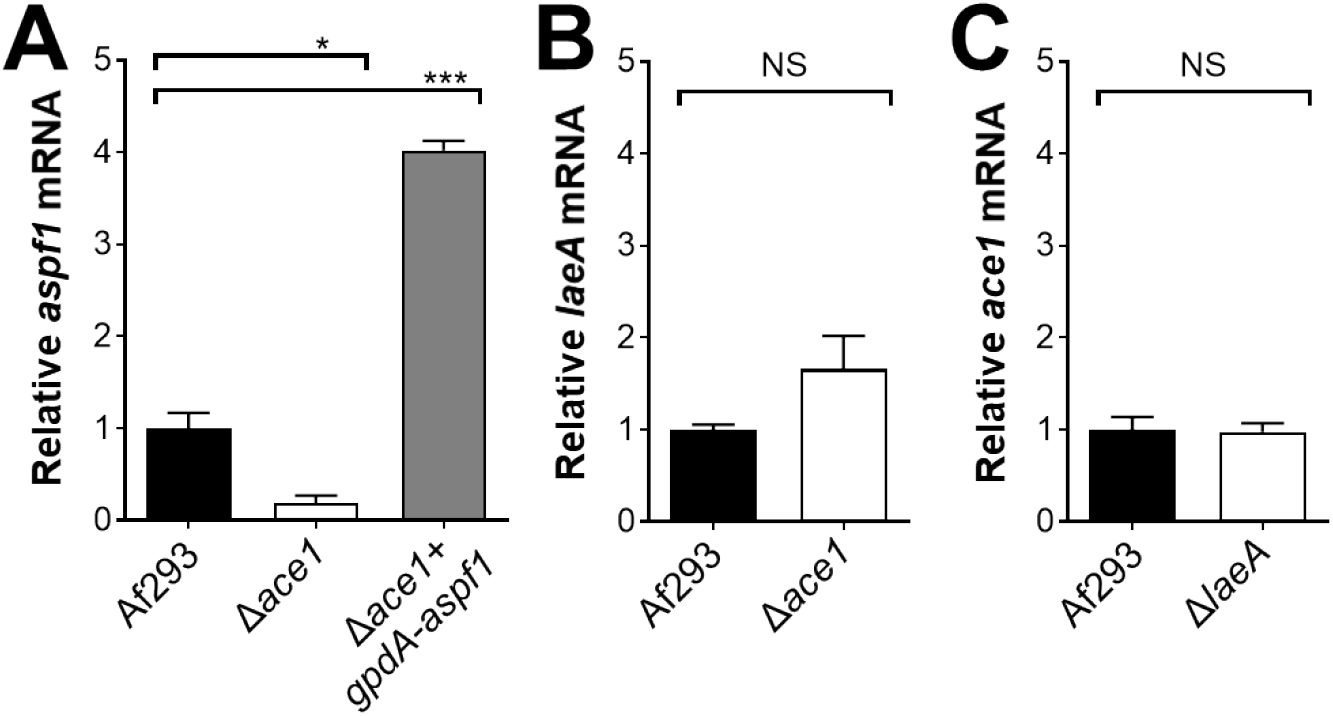
qPCR verification of transcriptional profiling results. Real-time PCR analysis of the relative transcript levels of *aspf1* (A), *laeA* (B), and *ace1* (C) in the indicates strains. The organisms were grown in AMM with low nitrogen and low zinc. Results are the mean ± SD of 3 biological replicates. *, *P* < 0.05; ***, *P* <0.001; NS, not significant.

A major regulator of secondary metabolite production in *A. fumigatus* is LaeA, which removes heterochromatic marks from the promoters of numerous genes, enabling the transcription of secondary metabolite genes [45, 46]. Microarray analysis indicates that LaeA governs the expression of at least 16 secondary metabolite gene clusters [45, 46]. Comparison of the microarray analysis of the Δ*laeA* mutant with the current RNA-seq analysis of the Δ*ace1* mutant indicates that Ace1 governs the expression of 6 biosynthetic gene clusters that are not known to be regulated by LaeA (Fig 6). LaeA and Ace1 differ in additional respects. All gene clusters that are regulated by LaeA are down-regulated in the Δl*aeA* mutant [46], whereas some gene clusters that are regulated by Ace1, such as fusarinine C, neosartoricin, and fumitremorgin, are up-regulated in the Δ*ace1* mutant. Also, LaeA governs asexual development and conidiation [45, 47], whereas we found no evidence that Ace1 governs these processes. These results suggest that Ace1 regulates the expression of secondary metabolite gene clusters by a different mechanism than LaeA.

To further investigate the relationship between Ace1 and LaeA, we used qPCR to measure the transcript levels of TF genes in the Δ*ace1* and Δ*laeA* mutant. The transcript levels of *laeA* were slightly higher in the Δ*ace1* mutant than in the wild-type strain, but this difference was not significant (Fig 7B). Also, *ace1* was expressed at wild-type levels in the Δ*laeA* mutant (Fig 7C). Overall, these data indicate that Ace1 regulates secondary metabolite gene clusters independently of LaeA.

### Ace1 governs virulence via Asp f1

The RNA-seq data suggested that the Δ*ace1* mutant had reduced virulence because of decreased production of mycotoxins. We investigated whether ergot alkaloids produced by the fumigaclavine biosynthesis cluster play a role in *A. fumigatus* virulence. The first enzyme in the fumigaclavine biosynthesis pathway is DmaW [48] and deletion of *dmaW* results in absent production of all detectable ergot alkaloids and attenuated virulence in *Galleria mellonella* [49]. We constructed a Δ*dmaW* mutant and analyzed its virulence in corticosteroid treated mice. The survival of mice infected with this mutant was similar to that of mice infected with the wild-type strain (Fig 8A), indicating that the reduced virulence of the Δ*ace1* mutant was not due to the absence of ergot alkaloid production.

**Fig 8.**
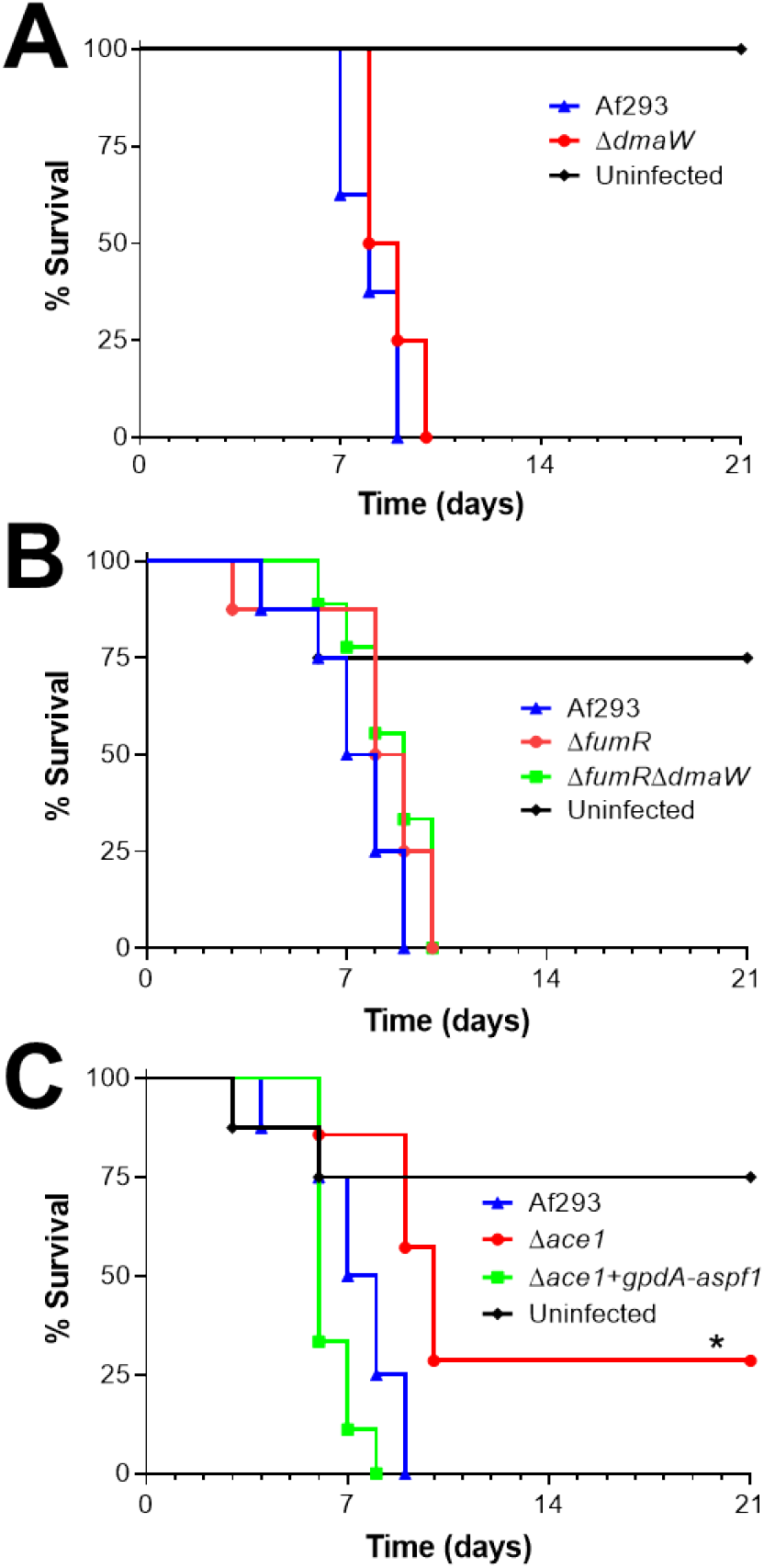
Forced expression of *aspf1* rescues the virulence defect of the Δ*ace1* mutant. Survival of mice infected with the indicated strains of *A. fumigatus*. Results are from 8 mice per strain. *, *p* < 0.

Another gene cluster that was down-regulated in the Δ*ace1* mutant was the large fumagillin and pseruotin supercluster. Fumagillin inhibits neutrophil function [50] and is required for *A. fumigatus* to cause maximal damage to the A549 pulmonary epithelial cell line [51]. Within the fumagillin biosynthetic gene cluster is *fumR*, which specifies a putative C6 type transcription factor that is required for fumagillin and pseurotin synthesis [38, 39]. We constructed a Δ*fumR* mutant and found that it had wild-type virulence in mice (Fig 8B). A Δ*fumR* Δ*dmaW* double mutant also had no detectable reduction in virulence (Fig 8B). Collectively, these data suggest that both fumagillin and the fumigaclavine ergot alkaloids are dispensable for virulence in the corticosteroid treated mouse model of invasive aspergillosis. Thus, the decreased production of these secondary metabolites does not explain the reduced virulence of the Δ*ace1* mutant.

Next, we investigated whether the attenuated virulence of the Δ*ace1* mutant was due to decreased expression of *aspf1,* which encodes a ribotoxin [37]. Previously, we have determined that Asp f1 is required for the maximal virulence of *A. fumigatus* in corticosteroid treated mice [12]. We constructed a variant of the Δ*ace1* mutant in which the expression of *aspf1* was driven by the constitutive *gpdA* promoter (Fig 7A). The forced expression of *aspf1* restored the virulence of the Δ*ace1* mutant to wild-type levels (Fig 8C). Thus, the reduced expression of *aspf1* likely accounts for the decreased virulence of the Δ*ace1* mutant.

Collectively, out results indicate that invasive growth in the lungs of corticosteroid treated mice induces a unique transcription profile in *A. fumigatus* as the organism responds to nutrient limitation and attack by host phagocytes. Also, growth in AMM with low zinc and low nitrogen *in vitro* induces a transcriptional response that largely mimics that induced by growth *in vivo*. This set of conditions can be used for RNA-seq analysis of *A. fumigatus* TF gene mutants to identify potential downstream target genes whose products mediate virulence. NanoString profiling of *A. fumigatus* during invasive growth in the lungs identified RlmA as a transcription factor that governs proliferation *in vivo*. It also identified Ace1 as a transcription factor that governs pathogenicity by regulating the expression of multiple secondary metabolite gene clusters and *aspf1* independently of LaeA. In our NanoString dataset are additional TF genes that are either up-regulated or highly expressed during invasive growth *in vivo*. Determining the roles of these genes in governing *A. fumigatus* virulence is currently in progress.

## Methods

### Ethics statement

All mouse studies were conducted in compliance with the NIH Guide for the Care and Use of Laboratory Animals. The experimental procedures were approved in advance by the Institutional Animal Care and Use Committee at the Lundquist Institute. The mice were group-housed according to experimental group in HEPA-filtered laminar flow cages with unrestricted access to food and water. The vivarium is managed by the Lundquist Institute in compliance with all policies and regulations of the Office of Laboratory Animal Welfare of the Public Health Service. The facility is fully accredited by the American Association for Laboratory Animal Care.

### Strains, media and growth conditions

The *A. fumigatus* strains used in this study are listed in Table 4. All strains were grown on Sabouraud dextrose agar (Difco) at 37°C for 7 d prior to use. Conidia were harvested with phosphate-buffered saline (PBS) containing 0.1% Tween 80 (Sigma-Aldrich) and enumerated with a hemacytometer.

**Table 4.** *A. fumigatus* strains used in the current study.

| Strain | Genotype | Reference |
| --- | --- | --- |
| Af293 | Wild-type | [67] |
| $\Delta rlmA$ | Af293; *rlmA::hph* | Present study |
| $\Delta ace1$ | Af293; *ace1::hph* | Present study |
| $\Delta afu1g15910$ | Af293; *Afu1g15910::hph* | Present study |
| $\Delta afu3g08050$ | Af293; *Afu3g08050::hph* | Present study |
| $\Delta afu4g10220$ | Af293; *Afu4g10220::hph* | Present study |
| $\Delta afu5g01650$ | Af293; *Afu5g01650::hph* | Present study |
| $\Delta afu5g13790$ | Af293; *Afu5g13790::hph* | Present study |
| $\Delta afu7g00130$ | Af293; *Afu7g00130::hph* | Present study |
| $\Delta afu8g05270$ | Af293; *Afu8g05270::hph* | Present study |
| $\Delta rlmA+rlmA$ | $\Delta rlmA$; *rlmA*; *ble* | Present study |
| $\Delta ace1+ace1$ | $\Delta ace1$; *ace1*; *ble* | Present study |
| $\Delta laeA$ | Af293.1; *laeA::pyrG1* | [46] |
| $\Delta dmaW$ | Af293; *dmaW::hph* | Present study |
| $\Delta fumR$ | Af293; *fumR::hph* | Present study |
| $\Delta fumR\Delta dmaW$ | $\Delta dmaW$; *fumR::ble* | Present study |
| $\Delta ace1+gpdA-aspf1$ | $\Delta ace1$; *gpdA-aspf1*; *ble* | Present study |
| Af293+*gpdA-GFP* | Af293; *gpdA-GFP*; *ble* | [55] |
| $\Delta ace1+gpdA-GFP$ | $\Delta ace1$; *gpdA-GFP*; *ble* | Present study |
| $\Delta ace1+ace1+gpdA-GFP$ | $\Delta ace1+ace1$; *gpdA-GFP*; *ptrA* | Present study |

For *in vitro* gene expression profiling analysis, 1.5 ×10^8^ conidia of *A. fumigatus* were added to 300 ml liquid standard AMM or modified AMM (no added iron, no added zinc, or the addition of 10% fetal bovine serum) and incubated for 24 h at 37°C in a shaking incubator. For growth under low nitrogen conditions, the *A. fumigatus* cells were grown in either AMM or modified AMM for 20 h, after which the hyphae were collected by filtration, washed with water, then added to AMM or modified AMM without nitrate and incubated for an additional 4 h. At the end of the incubation period, the resulting hyphae were collected by filtration and the RNA was extracted using the RNeasy Plant Minikit (Qiagen) following the manufacturer’s instructions.

For RNA-seq analysis, 1.5 × 10^8^ conidia of *A. fumigatus* were incubated in 300 ml of AMM without zinc for 20 h and then incubated in AMM without zinc and nitrate for 4 additional h prior to RNA extraction.

### Strain construction

The TF gene mutant strains used in this research were constructed using a split marker strategy. For each gene, approximately 1.5 kb of the 5’-flanking sequence upstream of the protein coding region was PCR-amplified from genomic DNA of strain Af293 using primers F3 and F4. The PCR primers used in the experiments are listed in Supplemental Table S2. The resulting fragment was cloned into plasmid pNLC106 [52]. Using the plasmid as a template, the 5’-flanking sequence region was linked to the 5’ portion of the *hph* hygromycin resistance gene by fusion PCR using primers F4 and HY. Next, about 1.5 kb of the 3’-flanking sequence downstream of the protein coding region was PCR-amplified from genomic DNA of strain A293 using primers F2 and F1 and the sequence of *hph* was amplified from pAN7 [53] using primers HYG-F and HYG-R. Using a mixture of the two fragments as the template, the 3’-flanking sequence region of target gene linked to 3’ portion of *hph* was amplified by fusion PCR with primers F1 and YG. Finally, the two fragments were used to transform Af293 protoplasts. The hygromycin-resistant clones were screened for the deletion of target gene by colony PCR using primers Screen-F and Screen-R.

To construct the Δ*rlmA* + *rlmA* complemented strain, a 4481 bp fragment containing the *rlmA* protein coding sequence and approximately 2 kb of 5’ flanking sequence and 0.5 kb of 3’ flanking sequence was PCR-amplified from Af293 genomic DNA using primers 3g08520-Com-F and 3g08520-Com-R. Similarly, to construct the Δ*ace1*+*ace1* complemented strain, a 4997 bp fragment containing the *ace1* protein coding sequence and flanking regions was PCR-amplified from using primers 3g08010-Com-F and 3g08010-Com-R. Each fragment was cloned into the NotI/XbaI sites of plasmid p402 [54]. The resulting plasmids were used to transform the Δ*rlmA* and Δ*ace1* strain. To confirm the presence of the complementation plasmids, the phleomycin-resistant colonies were screened by colony PCR using primers 3g08520-Com-F and 3g08520-Com-R to detect *rlmA* or 3g08010-Com-F and 3g08010-Com-R to detect *ace1*. The transcript levels of *rlmA* or *ace1* in various clones were quantified by real-time RT-PCR using primers RT-F and RT-R. The clones in which the transcript levels of *rlmA* or *ace1* was most similar to that of the wild-type strain was used in all experiments.

For use in the epithelial cell invasion assays, strains of *A. fumigatus* that expressed GFP were constructed. The Δ*ace1* mutant was transformed with plasmid GFP-Phleo and the Δ*ace1+ace1* complemented strain was transformed with plasmid GFP-pPTRI [55].

To construct the Δ*dmaW* (Afu2g18040) and Δ*fumR* (Afu8g00420) deletion mutants, a transient CRISPR-Cas9 gene deletion system was used [56, 57]. The Cas9 expression cassette was amplified from plasmid pFC331 [57], using primers Cas9-F and Cas9-R. To construct the sgRNA expression cassette, two DNA fragments were amplified from plasmid pFC334 [57] using primers sgRNA-F and sgRNA-ss-R, and sgRNA-R, sgRNA-ss-F. Next, the sgRNA expression cassette was amplified by fusion PCR from the two DNA fragments, using primers sgRNA-F and sgRNA-R. The hygromycin resistance (HygR) repair template was amplified from plasmid pVG2.2-hph [58] using primers Hyg-F and Hyg-R, which had about 50 bp of homology to the 5’ end of the protein coding sequence of the gene and the 3’ end of the protein coding sequence, respectively. The HygR repair template was mixed with the Cas9 cassette and the two sgRNA cassettes and then used for protoplast transformation. Hygromycin resistant clones were screened for deletion of target gene by colony PCR using primers RT-F and RT-R. The positive clones were also confirmed for absence of integration of DNA encoding Cas9 or the gRNA, using primers Cas9RT-F and Cas9RT-R, and sgRT-F, sgRT-R.

The Δ*fumR* Δ*dmaW* double mutant was constructed using the above CRISPR-Cas9 approach starting with the Δ*dmaW* mutant strain. The phleomycin (PhlR) repair template was amplified from plasmid p402 using primers Phleo-F and Phleo-R, which contained regions that were homologous to the 5’ and 3’ ends of the *fumR* protein coding sequence. Protoplasts were transformed with the repair template along with the Cas9 and sgRNA cassettes that were used to construct the Δ*fumR* deletion mutant. Phleomycin resistant clones were screened by colony PCR to identify ones with deletion of both genes and that lacked Cas9 and sgRNA sequences.

A strain of the Δ*ace1* mutant in which *aspf1* expression was driven by the *gpdA* promoter was constructed using the CRISPR-Cas9 system. The protein coding region of *aspf1* was amplified from genomic DNA using primers Aspf1-F and Aspf1-R. The resulting fragment was cloned into the *BamHI-NcoI* sites of plasmid pGFP-phleo using the NEBuilder DNA assembly kit (New England Biolabs). In this plasmid, the expression of *aspf1* was driven by the *A. nidulans gpdA* promoter. The *gpdA-aspf1*-phleomycin template was PCR amplified from this plasmid using primers Phleo-OE-F and Phleo-OE-R (Supplemental Table S1), which had about 50 bp of homology to the safe haven region of the *A. fumigatus* genome [59]. The Δ*ace1* mutant was transformed with the *gpdA-aspf1*-phleomycin template, the Cas9 cassette and the two safe haven sgRNA cassettes [59]. In the resulting phleomycin resistant clones, the *aspf1* transcript levels were quantified by real-time RT-PCR using primers Aspf1-RT-F and Aspf1-RT-R. A clone in which the *aspf1* mRNA expression was approximately 4-fold higher than the wild-type strain was used in all subsequent experiments.

### Mouse model of invasive pulmonary aspergillosis

A non-neutropenic, immunosuppressed mouse model of invasive aspergillosis was used to assess the transcriptional profile and virulence of the various strains [19]. Briefly, 6 week old, male Balb/c mice (Taconic Laboratories) were immunosuppressed with 7.5 mg cortisone acetate (Sigma-Aldrich) administered subcutaneously every other day starting at day −4 before infection for a total of 5 doses. To prevent bacterial infections, enrofloxacin (Baytril, Western Medical Supply) was added to the drinking water at a final concentration of 0.005% on day −5 relative to infection. The mice were infected by placing them for 1 h in an acrylic chamber into which 12 ml of 1×10^9^ conida/ml were aerosolized. Control mice were immunosuppressed, but not infected.

For the transcriptional profiling experiments, 3 mice infected with each strain were sacrificed after 2, 4, and 5 days infection. Their lungs were harvested and snap frozen in liquid nitrogen for RNA extraction. To isolate fungal RNA from the infected mouse lungs, the RNeasy minikit (Qiagen) was used with modifications [12]. Approximately 2.4 ml of buffer RLT with 1% β-mercaptoethanol was added to the lungs from each mouse and the tissue was homogenized in an M tube (Miltenyi Biotec) using a gentleMACS dissociator (Miltenyi Biotec) on setting RNA_02.01. Next, the homogenate was mixed with an equal volume of phenol-chloroform-isoamyl alcohol (25:24:1) and a half volume of zirconium beads (Ambion) and then vortexed with a Mini-Beadbeater (Biospec Products) for 3 min. After centrifugation, the aqueous phase was collected and mixed with an equal volume of 70% ethanol. The RNA was isolated from this mixture using an RNeasy spin column (Qiagen) following the manufacturer’s instructions.

To assess the virulence of the various *A. fumigatus* strains using survival as the end point, 11 mice were infected with each strain. Shortly after infection, 3 mice from each group were sacrificed, and their lungs were harvested, homogenized and quantitatively cultured to verify conidia delivery to the lung. The remaining mice were monitored twice daily for survival. 5 mice that were immunosuppressed, but not infected were included as a negative control.

### NanoString analysis

A NanoString nCounter digital analyzer was used for transcriptional profiling of *A. fumigatus* Af293 both *in vivo* and *in vitro* as previously described [12]. For each condition, the adjusted data were normalized to total probe counts. However, when we compared 18 genes that were in common between our two probe sets (called TF and ER), we observed poor agreement in fold changes. The ER genes had been chosen because they respond dramatically to environmental changes, and we reasoned that large expression changes may make total counts unreliable for normalization. The TF probe set, representing 400 different putative transcription factor genes, is extremely diverse and thus total counts are more reliable for normalization. With that point in mind, the ER datasets were renormalized to TF dataset measurements as follows. For each of the 18 common genes in both ER and TF probe sets, we calculated the ratio of mean TF counts/mean ER counts for each growth condition. Then the median ratio for the 18 genes was used to calculate a normalization factor for each growth condition. The normalization factor for each growth condition was applied to all ER genes. Finally, the TF and ER datasets were combined, with counts for common genes from the TF datasets, and 10 TF genes with the lowest counts removed. Each condition was tested in 3 biological replicates and the expression ratios were calculated using the mean values. Genes were considered differentially expressed when there was at least a 2-fold change in the transcript levels and an unpaired, two-tailed student’s t-test p-value ≤ 0.05.

### RNA-seq

RNA-seq libraries (strand-agnostic, 150 bp paired-end) were generated from total fungal RNA by Novogene Corporation Inc. Sequencing reads were aligned to the reference *A*. *fumigatus* Af293 genome using HISAT2 [60] and alignment files were used to generate read counts for each gene using HTseq [61]. We obtained an average of 46.4 million aligned reads per sample. Statistical analysis of differential gene expression was performed using the DEseq package from Bioconductor [62]. A gene was considered differentially expressed if the FDR value for differential expression was below 0.05. The RNA-seq analysis was performed in biological triplicate. The raw RNA-seq data will be deposited at the NCBI Short Read Archive (SRA) data base.

### Stress assays

To test the susceptibility of the various strains to cell wall, cell membrane, oxidant, and ionic stress, serial 10-fold dilutions of conidia ranging from 10^5^ to 10^2^ cells in a volume of 5 μl were spotted onto AMM agar plates supplemented with 5 mM protamine (Sigma-Aldrich), 0.01% SDS (Sigma), 40 μg/ml caspofungin (Bellavida Pharmacy), 200 ug/ml Congo red (Sigma-Aldrich), 300 μg/ml Calcofluor White (Sigma-Aldrich), 4 mM H_2_O_2_, 200 mM KCl, 200 mM MgCl_2_, 200 mM NaCl, or 50 mM CaCl_2_. To determine growth at alkaline pH, the conidia were spotted onto AMM agar adjusted to pH 8.0 with NaOH. Fungal growth was analyzed after incubation at 37°C for 2 d.

### Real-time PCR

The total RNA was reverse transcribed into cDNA using Moloney murine leukemia virus reverse transcriptase (Promega). Real-time PCR was performed using the POWER SYBR green PCR master mix (Applied Biosystems) and an ABI 7000 thermocycler (Applied Biosystems). Gene transcript levels were quantified by ΔΔC^t^ method, using GAPDH as the endogenously expressed gene [63].

### Host cell interaction assays

The capacity of the various strains to adhere to, invade, and damage the A549 pulmonary epithelial cell line (American Type Culture Collection) was determined using our previously described methods [11, 55, 64]. To measure adherence and invasion, 10^5^ germlings of the various GFP-expressing strains of *A. fumigatus* in F12k medium (American Type Culture Collection) were added to A549 cells that had been grown to confluency in 24-well tissue culture plates containing fibronectin coated circular glass coverslips in each well. After incubation for 2.5 h, the cells were rinsed with 1 ml HBSS in a standardized manner and then fixed with 3% paraformaldehyde. The noninternalized portions of the organisms were stained with a polyclonal rabbit anti-*A. fumigatus* primary antibody (Meridian Life Science, Inc.) followed by an AlexaFluor 568-labeled secondary antibody (Life Technologies). After the coverslips were mounted inverted on microscope slides, they were viewed by epifluorescence. The number of cell-associated organisms was determined by counting the number of GFP-expressing organisms per high-powered field (HPF). The number of endocytosed organisms was determined by subtracting the number of non-internalized organisms (which fluoresced red) from the number of cell-associated organisms. At least 100 organisms per coverslip were scored and each strain was tested in triplicate in three independent experiments.

Our standard ^51^Cr release assay was used to evaluate the capacity of the various strains to damage the A549 cell line [11, 65]. The A549 cells were grown to confluency in a 24-well tissue culture plate and then loaded with ^51^Cr overnight. After rinsing the cells to remove the unincorporated ^51^Cr, the cells were infected with 5×10^5^ conidia of each strain in F12K medium. After 16 h of infection, the medium above the cells was collected and the cells were lysed with 6 N NaOH. The lysed cells were collected by rinsing the wells twice with RadiacWash (Biodex Medical Systems). The amount of ^51^Cr in the medium and the cell lysate was measured using a gamma counter. The spontaneous release of ^51^Cr was determined using uninfected A549 cells that were processed in parallel. The specific release of ^51^Cr was calculated using our previous described formula [11, 65]. Each experiment was performed in triplicate and repeated three times.

The susceptibility of the various *A. fumigatus* strains to phagocyte killing was determined using bone marrow-derived macrophages (BMDMs), which were isolated from 6-week-old mice (Taconic Laboratories). The cells were differentiated into macrophages by incubation with 50 ng/ml macrophage colony–stimulating factor (M-CSF) (BioLegend) in Dulbecco’s Modified Eagle’s Medium (DMEM) (American Type Culture Collection) with 10% fetal bovine serum (Gemini Bio-Products), 1% streptomycin and penicillin for 10 d [66]. The day before the experiment, the adherent cells were harvested and 10^6^ cells were seeded into each well of a 6-well tissue culture plate. The next day, 5 × 10^4^ conidia were added to each well and incubated for 8 h. A similar number of conidia was added to a second 6-well tissue culture plate without BMDMs as a control. At the end of the incubation period, the BMDMs were lysed with distilled water and sonication. The contents of the wells were aspirated and quantitatively cultured on Sabouraud dextrose agar. For each strain, the percentage of *A. fumigatus* cells killed was calculated by the formula: 1 - number of colonies in the wells containing BMDMs/ number of colonies in the wells without BMDMs. Each experiment was performed in triplicates and repeated three times.

### Statistical analysis

The data from the in vitro experiments were analyzed by the two-tailed Student’s t-test assuming unequal variance or one way analysis of variance followed by the Dunnett’s test for multiple comparisons. The survival data were analyzed using the Log-Rank test. A *P*-value of ≤ 0.05 was considered to be significant.

## ACKNOWLEDGEMENTS

We thank the members of the Filler, Bruno, and Mitchell labs for helpful discussions and suggestions.

## REFERENCES

1. Dagenais TR, Keller NP. Pathogenesis of *Aspergillus fumigatus* in invasive aspergillosis. Clin Microbiol Rev. 2009;22(3):447–65. Epub 2009/07/15. doi: 22/3/447 [pii] 10.1128/CMR.00055-08. PubMed PMID: 19597008; PubMed Central PMCID: PMC2708386.

2. Segal BH. Aspergillosis. N Engl J Med. 2009;360(18):1870–84. Epub 2009/05/01. doi: 360/18/1870 [pii] 10.1056/NEJMra0808853. PubMed PMID: 19403905.

3. Sherif R, Segal BH. Pulmonary aspergillosis: clinical presentation, diagnostic tests, management and complications. Curr Opin Pulm Med. 2010;16(3):242–50. Epub 2010/04/09. doi: 10.1097/MCP.0b013e328337d6de 00063198-201005000-00012 [pii]. PubMed PMID: 20375786; PubMed Central PMCID: PMC3326383.

4. Walsh TJ, Anaissie EJ, Denning DW, Herbrecht R, Kontoyiannis DP, Marr KA, et al. Treatment of aspergillosis: clinical practice guidelines of the Infectious Diseases Society of America. Clin Infect Dis. 2008;46(3):327–60. Epub 2008/01/08. doi: 10.1086/525258. PubMed PMID: 18177225.

5. Georgiadou SP, Kontoyiannis DP. The impact of azole resistance on aspergillosis guidelines. Ann N Y Acad Sci. 2012;1272:15–22. Epub 2012/12/13. doi: 10.1111/j.1749-6632.2012.06795.x. PubMed PMID: 23231710.

6. Pfaller MA. Antifungal drug resistance: mechanisms, epidemiology, and consequences for treatment. Am J Med. 2012;125(1 Suppl):S3–13. Epub 2012/01/04. doi: S0002-9343(11)00913-2 [pii] 10.1016/j.amjmed.2011.11.001. PubMed PMID: 22196207.

7. Bertuzzi M, Schrettl M, Alcazar-Fuoli L, Cairns TC, Munoz A, Walker LA, et al. The pH-responsive PacC transcription factor of *Aspergillus fumigatus* governs epithelial entry and tissue invasion during pulmonary aspergillosis. PLoS Pathog. 2014;10(10):e1004413. Epub 2014/10/21. doi: 10.1371/journal.ppat.1004413. PubMed PMID: 25329394; PubMed Central PMCID: PMCPMC4199764.

8. McDonagh A, Fedorova ND, Crabtree J, Yu Y, Kim S, Chen D, et al. Sub-telomere directed gene expression during initiation of invasive aspergillosis. PLoS Pathog. 2008;4(9):e1000154. PubMed PMID: 18787699.

9. Kale SD, Ayubi T, Chung D, Tubau-Juni N, Leber A, Dang HX, et al. Modulation of immune signaling and metabolism highlights host and fungal transcriptional responses in mouse models of invasive pulmonary aspergillosis. Sci Rep. 2017;7(1):17096. Epub 2017/12/08. doi: 10.1038/s41598-017-17000-1. PubMed PMID: 29213115; PubMed Central PMCID: PMCPMC5719083.

10. Abad A, Fernandez-Molina JV, Bikandi J, Ramirez A, Margareto J, Sendino J, et al. What makes *Aspergillus fumigatus* a successful pathogen? Genes and molecules involved in invasive aspergillosis. Rev Iberoam Micol. 2010;27(4):155–82.

11. Pongpom M, Liu H, Xu W, Snarr BD, Sheppard DC, Mitchell AP, et al. Divergent targets of *Aspergillus fumigatus* AcuK and AcuM transcription factors during growth in vitro versus invasive disease. Infect Immun. 2015;83(3):923–33.

12. Liu H, Xu W, Solis NV, Woolford C, Mitchell AP, Filler SG. Functional convergence of gliP and aspf1 in *Aspergillus fumigatus* pathogenicity. Virulence. 2018;9(1):1062–73.

13. Xu W, Solis NV, Ehrlich RL, Woolford CA, Filler SG, Mitchell AP. Activation and alliance of regulatory pathways in *C. albicans* during mammalian infection. PLoS Biol. 2015;13(2):e1002076.

14. Fanning S, Xu W, Solis N, Woolford CA, Filler SG, Mitchell AP. Divergent targets of *Candida albicans* biofilm regulator Bcr1 in vitro and in vivo. Eukaryot Cell. 2012;11:896–904. Epub 2012/05/01. doi: 10.1128/EC.00103-12. PubMed PMID: 22544909.

15. Nobile CJ, Solis N, Myers CL, Fay AJ, Deneault JS, Nantel A, et al. *Candida albicans* transcription factor Rim101 mediates pathogenic interactions through cell wall functions. Cell Microbiol. 2008;10(11):2180–96. PubMed PMID: 18627379.

16. Nobile CJ, Andes DR, Nett JE, Smith FJ, Yue F, Phan QT, et al. Critical role of Bcr1-dependent adhesins in *C. albicans* biofilm formation in vitro and in vivo. PLoS Pathog. 2006;2(7):e63. PubMed PMID: 16839200.

17. Davis DA, Bruno V, Loza L, Filler SG, Mitchell AP. *C. albicans* Mds3p, a conserved regulator of pH responses and virulence identified through insertional mutagenesis. Genetics. 2002;162(4):1573–81. PubMed PMID: 12524333.

18. Bahn YS. Exploiting fungal virulence-regulating transcription factors as novel antifungal drug targets. PLoS Pathog. 2015;11(7):e1004936. Epub 2015/07/17. doi: 10.1371/journal.ppat.1004936. PubMed PMID: 26181382; PubMed Central PMCID: PMC4504714.

19. Chiang LY, Sheppard DC, Gravelat FN, Patterson TF, Filler SG. *Aspergillus fumigatus* stimulates leukocyte adhesion molecules and cytokine production by endothelial cells in vitro and during invasive pulmonary disease. Infect Immun. 2008;76(8):3429–38. PubMed PMID: 18490455.

20. Sheppard DC, Rieg G, Chiang LY, Filler SG, Edwards JE, Jr., Ibrahim AS. Novel inhalational murine model of invasive pulmonary aspergillosis. Antimicrob Agents Chemother. 2004;48(5):1908–11. PubMed PMID: 15105158.

21. Schrettl M, Kim HS, Eisendle M, Kragl C, Nierman WC, Heinekamp T, et al. SreA-mediated iron regulation in Aspergillus fumigatus. Mol Microbiol. 2008;70(1):27–43. PubMed PMID: 18721228.

22. Schrettl M, Beckmann N, Varga J, Heinekamp T, Jacobsen ID, Jochl C, et al. HapX-mediated adaption to iron starvation is crucial for virulence of *Aspergillus fumigatus*. PLoS Pathog. 2010;6(9):1001124.

23. Schrettl M, Bignell E, Kragl C, Joechl C, Rogers T, Arst HN, Jr., et al. Siderophore biosynthesis but not reductive iron assimilation is essential for *Aspergillus fumigatus* virulence. J Exp Med. 2004;200(9):1213–9.

24. Moreno MA, Ibrahim-Granet O, Vicentefranqueira R, Amich J, Ave P, Leal F, et al. The regulation of zinc homeostasis by the ZafA transcriptional activator is essential for *Aspergillus fumigatus* virulence. Mol Microbiol. 2007;64(5):1182–97. Epub 2007/06/05. doi: 10.1111/j.1365-2958.2007.05726.x. PubMed PMID: 17542914.

25. Hensel M, Arst HN, Jr., Aufauvre-Brown A, Holden DW. The role of the *Aspergillus fumigatus* areA gene in invasive pulmonary aspergillosis. Mol Gen Genet. 1998;258(5):553–7. Epub 1998/07/21. PubMed PMID: 9669338.

26. Willger SD, Puttikamonkul S, Kim KH, Burritt JB, Grahl N, Metzler LJ, et al. A sterol-regulatory element binding protein is required for cell polarity, hypoxia adaptation, azole drug resistance, and virulence in *Aspergillus fumigatus*. PLoS Pathog. 2008;4(11):e1000200. PubMed PMID: 18989462.

27. Richie DL, Hartl L, Aimanianda V, Winters MS, Fuller KK, Miley MD, et al. A role for the unfolded protein response (UPR) in virulence and antifungal susceptibility in *Aspergillus fumigatus*. PLoS Pathog. 2009;5(1):e1000258. PubMed PMID: 19132084.

28. Dinamarco TM, Almeida RS, de Castro PA, Brown NA, dos Reis TF, Ramalho LN, et al. Molecular characterization of the putative transcription factor SebA involved in virulence in Aspergillus fumigatus. Eukaryot Cell. 2012;11(4):518–31.

29. Dirr F, Echtenacher B, Heesemann J, Hoffmann P, Ebel F, Wagener J. AfMkk2 is required for cell wall integrity signaling, adhesion, and full virulence of the human pathogen *Aspergillus fumigatus*. Int J Med Microbiol. 2010;300(7):496–502.

30. Ma Y, Qiao J, Liu W, Wan Z, Wang X, Calderone R, et al. The sho1 sensor regulates growth, morphology, and oxidant adaptation in *Aspergillus fumigatus* but is not essential for development of invasive pulmonary aspergillosis. Infect Immun. 2008;76(4):1695–701.

31. Spikes S, Xu R, Nguyen CK, Chamilos G, Kontoyiannis DP, Jacobson RH, et al. Gliotoxin production in *Aspergillus fumigatus* contributes to host-specific differences in virulence. J Infect Dis. 2008;197(3):479–86. PubMed PMID: 18199036.

32. Balibar CJ, Walsh CT. GliP, a multimodular nonribosomal peptide synthetase in *Aspergillus fumigatus*, makes the diketopiperazine scaffold of gliotoxin. Biochemistry. 2006;45(50):15029–38.

33. Scharf DH, Remme N, Habel A, Chankhamjon P, Scherlach K, Heinekamp T, et al. A dedicated glutathione S-transferase mediates carbon-sulfur bond formation in gliotoxin biosynthesis. J Am Chem Soc. 2011;133(32):12322–5.

34. Bok JW, Chung D, Balajee SA, Marr KA, Andes D, Nielsen KF, et al. GliZ, a transcriptional regulator of gliotoxin biosynthesis, contributes to *Aspergillus fumigatus* virulence. Infect Immun. 2006;74(12):6761–8.

35. Smith TD, Calvo AM. The mtfA transcription factor gene controls morphogenesis, gliotoxin production, and virulence in the opportunistic human pathogen *Aspergillus fumigatus*. Eukaryot Cell. 2014;13(6):766–75.

36. Chooi YH, Fang J, Liu H, Filler SG, Wang P, Tang Y. Genome mining of a prenylated and immunosuppressive polyketide from pathogenic fungi. Org Lett. 2013;15(4):780–3.

37. Arruda LK, Platts-Mills TA, Fox JW, Chapman MD. *Aspergillus fumigatus* allergen I, a major IgE-binding protein, is a member of the mitogillin family of cytotoxins. J Exp Med. 1990;172(5):1529–32.

38. Dhingra S, Lind AL, Lin HC, Tang Y, Rokas A, Calvo AM. The fumagillin gene cluster, an example of hundreds of genes under veA control in *Aspergillus fumigatus*. PLoS ONE. 2013;8(10).

39. Wiemann P, Guo CJ, Palmer JM, Sekonyela R, Wang CC, Keller NP. Prototype of an intertwined secondary-metabolite supercluster. Proc Natl Acad Sci U S A. 2013;110(42):17065–70.

40. Yin WB, Baccile JA, Bok JW, Chen Y, Keller NP, Schroeder FC. A nonribosomal peptide synthetase-derived iron(III) complex from the pathogenic fungus *Aspergillus fumigatus*. J Am Chem Soc. 2013;135(6):2064–7.

41. Throckmorton K, Lim FY, Kontoyiannis DP, Zheng W, Keller NP. Redundant synthesis of a conidial polyketide by two distinct secondary metabolite clusters in *Aspergillus fumigatus*. Environ Microbiol. 2016;18(1):246–59. Epub 2015/08/06. doi: 10.1111/1462-2920.13007. PubMed PMID: 26242966; PubMed Central PMCID: PMCPMC4750049.

42. Rocha MC, Fabri JH, Franco de Godoy K, Alves de Castro P, Hori JI, Ferreira da Cunha A, et al. *Aspergillus fumigatus* MADS-Box Transcription Factor rlmA Is Required for Regulation of the Cell Wall Integrity and Virulence. G3 (Bethesda). 2016;6(9):2983–3002. Epub 2016/07/31. doi: 10.1534/g3.116.031112. PubMed PMID: 27473315; PubMed Central PMCID: PMCPMC5015955.

43. Shantappa S, Dhingra S, Hernandez-Ortiz P, Espeso EA, Calvo AM. Role of the zinc finger transcription factor SltA in morphogenesis and sterigmatocystin biosynthesis in the fungus *Aspergillus nidulans*. PLoS One. 2013;8(7):e68492. Epub 2013/07/11. doi: 10.1371/journal.pone.0068492. PubMed PMID: 23840895; PubMed Central PMCID: PMCPMC3698166.

44. Spielvogel A, Findon H, Arst HN, Araujo-Bazan L, Hernandez-Ortiz P, Stahl U, et al. Two zinc finger transcription factors, CrzA and SltA, are involved in cation homoeostasis and detoxification in *Aspergillus nidulans*. Biochem J. 2008;414(3):419–29. Epub 2008/05/13. doi: 10.1042/bj20080344. PubMed PMID: 18471095.

45. Lind AL, Lim FY, Soukup AA, Keller NP, Rokas A. An LaeA- and BrlA-dependent cellular network governs tissue-specific secondary metabolism in the human pathogen *Aspergillus fumigatus*. mSphere. 2018;3(2). Epub 2018/03/23. doi: 10.1128/mSphere.00050-18. PubMed PMID: 29564395; PubMed Central PMCID: PMCPMC5853485.

46. Perrin RM, Fedorova ND, Bok JW, Cramer RA, Wortman JR, Kim HS, et al. Transcriptional regulation of chemical diversity in *Aspergillus fumigatus* by LaeA. PLoS Pathog. 2007;3(4):e50. PubMed PMID: 17432932.

47. Bok JW, Balajee SA, Marr KA, Andes D, Nielsen KF, Frisvad JC, et al. LaeA, a regulator of morphogenetic fungal virulence factors. Eukaryot Cell. 2005;4(9):1574–82. PubMed PMID: 16151250.

48. Coyle CM, Panaccione DG. An ergot alkaloid biosynthesis gene and clustered hypothetical genes from *Aspergillus fumigatus*. Appl Environ Microbiol. 2005;71(6):3112–8. Epub 2005/06/04. doi: 10.1128/aem.71.6.3112-3118.2005. PubMed PMID: 15933009; PubMed Central PMCID: PMCPMC1151871.

49. Panaccione DG, Arnold SL. Ergot alkaloids contribute to virulence in an insect model of invasive aspergillosis. Sci Rep. 2017;7(1):8930. Epub 2017/08/23. doi: 10.1038/s41598-017-09107-2. PubMed PMID: 28827626; PubMed Central PMCID: PMCPMC5567044.

50. Fallon JP, Reeves EP, Kavanagh K. Inhibition of neutrophil function following exposure to the *Aspergillus fumigatus* toxin fumagillin. J Med Microbiol. 2010;59(Pt 6):625–33. Epub 2010/03/06. doi: 10.1099/jmm.0.018192-0. PubMed PMID: 20203215.

51. Guruceaga X, Ezpeleta G, Mayayo E, Sueiro-Olivares M, Abad-Diaz-De-Cerio A, Aguirre Urizar JM, et al. A possible role for fumagillin in cellular damage during host infection by *Aspergillus fumigatus*. Virulence. 2018;9(1):1548–61. Epub 2018/09/27. doi: 10.1080/21505594.2018.1526528. PubMed PMID: 30251593; PubMed Central PMCID: PMCPMC6177242.

52. Catlett NL, Lee B-N, Yoder OC, Turgeon BG. Split-marker recombination for efficient targeted deletion of fungal genes. Fungal Genet News. 2002;50:9–11.

53. Punt PJ, Oliver RP, Dingemanse MA, Pouwels PH, van den Hondel CA. Transformation of *Aspergillus* based on the hygromycin B resistance marker from *Escherichia coli*. Gene. 1987;56(1):117–24. PubMed PMID: 2824287.

54. Richie DL, Miley MD, Bhabhra R, Robson GD, Rhodes JC, Askew DS. The *Aspergillus fumigatus* metacaspases CasA and CasB facilitate growth under conditions of endoplasmic reticulum stress. Mol Microbiol. 2007;63(2):591–604. PubMed PMID: 17176258.

55. Liu H, Lee MJ, Solis NV, Phan QT, Swidergall M, Ralph B, et al. *Aspergillus fumigatus* CalA binds to integrin a5b1 and mediates host cell invasion. Nat Microbiol. 2016;2:16211.

56. Min K, Ichikawa Y, Woolford CA, Mitchell AP. *Candida albicans* gene deletion with a transient CRISPR-Cas9 system. mSphere. 2016;1(3). Epub 2016/06/25. doi: 10.1128/mSphere.00130-16. PubMed PMID: 27340698; PubMed Central PMCID: PMCPMC4911798.

57. Nødvig CS, Nielsen JB, Kogle ME, Mortensen UH. A CRISPR-Cas9 system for genetic engineering of filamentous fungi. PLoS One. 2015;10(7):e0133085. Epub 2015/07/16. doi: 10.1371/journal.pone.0133085. PubMed PMID: 26177455; PubMed Central PMCID: PMCPMC4503723.

58. Macheleidt J, Scherlach K, Neuwirth T, Schmidt-Heck W, Straßburger M, Spraker J, et al. Transcriptome analysis of cyclic AMP-dependent protein kinase A-regulated genes reveals the production of the novel natural compound fumipyrrole by *Aspergillus fumigatus*. Mol Microbiol. 2015;96(1):148–62. Epub 2015/01/15. doi: 10.1111/mmi.12926. PubMed PMID: 25582336; PubMed Central PMCID: PMCPMC4425693.

59. Pham T, Xie X, Lin X. An intergenic “safe haven” region in *Aspergillus fumigatus*. Med Mycol. 2020. Epub 2020/03/15. doi: 10.1093/mmy/myaa009. PubMed PMID: 32171003.

60. Kim D, Paggi JM, Park C, Bennett C, Salzberg SL. Graph-based genome alignment and genotyping with HISAT2 and HISAT-genotype. Nat Biotechnol. 2019;37(8):907–15. Epub 2019/08/04. doi: 10.1038/s41587-019-0201-4. PubMed PMID: 31375807; PubMed Central PMCID: PMCPMC7605509.

61. Anders S, Pyl PT, Huber W. HTSeq--a Python framework to work with high-throughput sequencing data. Bioinformatics. 2015;31(2):166–9. Epub 2014/09/28. doi: 10.1093/bioinformatics/btu638. PubMed PMID: 25260700; PubMed Central PMCID: PMCPMC4287950.

62. Anders S, Huber W. Differential expression analysis for sequence count data. Genome Biol. 2010;11(10):R106. Epub 2010/10/29. doi: 10.1186/gb-2010-11-10-r106. PubMed PMID: 20979621; PubMed Central PMCID: PMCPMC3218662.

63. Liu H, Gravelat FN, Chiang LY, Chen D, Vanier G, Ejzykowicz DE, et al. *Aspergillus fumigatus* AcuM regulates both iron acquisition and gluconeogenesis. Mol Microbiol. 2010;78(4):1038–54. Epub 2010/11/11. doi: 10.1111/j.1365-2958.2010.07389.x. PubMed PMID: 21062375; PubMed Central PMCID: PMC3051834.

64. Ejzykowicz DE, Cunha MM, Rozental S, Solis NV, Gravelat FN, Sheppard DC, et al. The *Aspergillus fumigatus* transcription factor Ace2 governs pigment production, conidiation and virulence. Mol Microbiol. 2009;72(1):155–69. PubMed PMID: 19220748.

65. Ejzykowicz DE, Solis NV, Gravelat FN, Chabot J, Li X, Sheppard DC, et al. Role of *Aspergillus fumigatus* DvrA in host cell interactions and virulence. Eukaryot Cell. 2010;9(10):1432–40. Epub 2010/08/03. doi: EC.00055-10 [pii] 10.1128/EC.00055-10. PubMed PMID: 20675576; PubMed Central PMCID: PMC2950423.

66. Zhang X, Goncalves R, Mosser DM. The isolation and characterization of murine macrophages. Curr Protoc Immunol. 2008;Chapter 14:Unit 14 1. Epub 2008/11/20. doi: 10.1002/0471142735.im1401s83. PubMed PMID: 19016445; PubMed Central PMCID: PMCPMC2834554.

67. Nierman WC, Pain A, Anderson MJ, Wortman JR, Kim HS, Arroyo J, et al. Genomic sequence of the pathogenic and allergenic filamentous fungus *Aspergillus fumigatus*. Nature. 2005;438(7071):1151–6. PubMed PMID: 16372009.

